# Modulation of Allostery with Multiple Mechanisms by Hotspot Mutations in TetR

**DOI:** 10.1101/2023.08.29.555381

**Authors:** Jiahua Deng, Yuchen Yuan, Qiang Cui

**Affiliations:** Department of Chemistry, Boston University, 590 Commonwealth Avenue, Boston, Massachusetts 02215, United States; Department of Physics, Boston University, 590 Commonwealth Avenue, Boston, Massachusetts 02215, United States; Department of Biomedical Engineering, Boston University, 44 Cummington Mall, Boston, Massachusetts 02215, United States

**Author notes:** J. D. and Y. Y. contributed equally to this work.

## Abstract

Modulating allosteric coupling offers unique opportunities for biomedical applications. Such efforts can benefit from efficient prediction and evaluation of allostery hotspot residues that dictate the degree of co-operativity between distant sites. We demonstrate that effects of allostery hotspot mutations can be evaluated qualitatively and semi-quantitatively by molecular dynamics simulations in a bacterial tetracycline repressor (TetR). The simulations recapitulate the effects of these mutations on abolishing the induction function of TetR and provide a rationale for the different degrees of rescuability observed to restore allosteric coupling of the hotspot mutations. We demonstrate that the same non-inducible phenotype could be the result of perturbations in distinct structural and energetic properties of TetR. Our work underscore the value of explicitly computing the functional free energy landscapes to effectively evaluate and rank hotspot mutations despite the prevalence of compensatory interactions, and therefore provide quantitative guidance to allostery modulation for therapeutic and engineering applications.

TOC Graphic

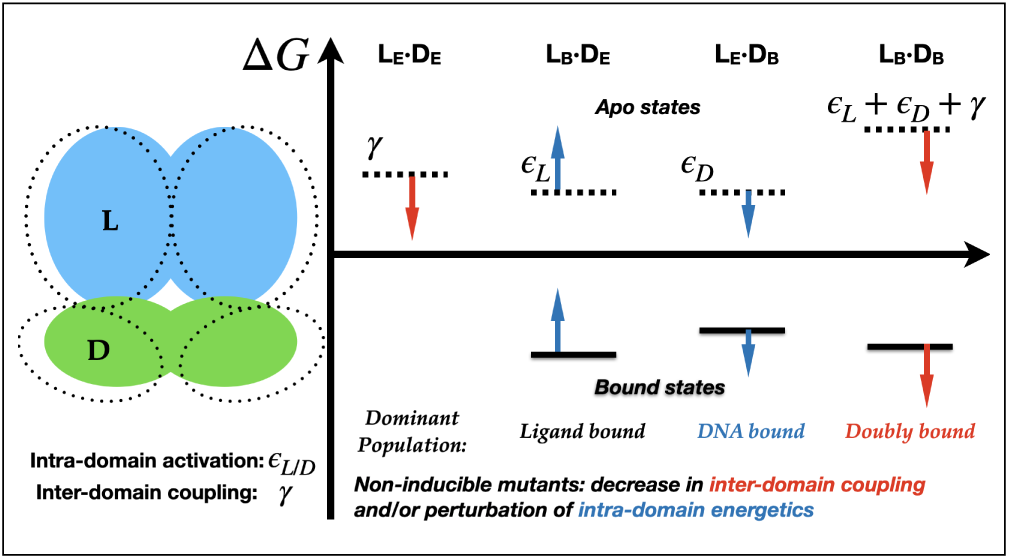

## Introduction

Allostery couples distant sites by binding an allosteric effector at the allosteric site, and propagation of structural or dynamical perturbations to the functional site to modulate its activity. It is essential to the function of proteins and nucleic acids in many biological processes.^1^ Controlling the allosteric coupling and communications is of broad therapeutic value and offers unique opportunities^2^ for biomedical and biotechnological applications. Designing molecules that target an allosteric site rather than the active site can potentially lead to higher levels of selectivity and specificity, especially for drug targets that feature highly conserved binding pockets such as kinases.^3,4^ Numerous serious human diseases such as various cancers are caused by mutations that either disrupt or activate allosteric communications,^5^ and resistance to drugs also often arises due to mutations that perturb allostery in the target protein.^6^ The understanding of allosteric communications also makes it feasible to design and engineer proteins with desired purposes by mutations, such as controlling enzyme activities for catalysis, constructing regulatory systems for gene expression, and developing biosensors and biological logic gates.^7–13^

To effectively design allosteric therapeutics and engineer/re-engineer allosteric proteins, it is useful to first identify and target allostery hotspots. Hotspots are the set of key residues that dictate the co-operativity between the effector site and the active site. In addition to their locations, the specific sequence features of allostery hotspot residues may also be important to their functional significances, since their contributions are likely sensitive to the local steric or polar interactions they are engaged in. Allostery hotspots can be explored by various computational techniques, such as statistical analysis of residual co-evolution,^14–16^ motional network and community analysis,^17–22^ mapping propagation of conformational distortions,^23^ and perturbation analysis of collective normal modes.^24–26^ Experimentally, site-directed mutagenesis,^27^ solution NMR chemical shift perturbations and relaxation measurements,^28–30^ surface plasmon resonance,^31^ and *in vitro* and *in vivo* biochemical essays/kinetics^32^ are commonly used approaches to probe allostery hotspots. As each methodology has its own limitation(s) in either cost or accuracy, it is still challenging to identify a complete set of allosteric hotspots, which prevents the evaluation of the individual role of each hotspot and a clear understanding of their mechanisms. A high-throughput methodology which could exhaustively and systematically analyze the sequence-function relationship is of great promise. Deep mutational scanning (DMS) analysis^33–35^ has emerged as a powerful technique for functional screening of allostery hotspots in proteins in a comprehensive and unbiased fashion. As demonstrated by recent applications,^36–41^ DMS analysis readily reveals both the location and sequence variability of allostery hotspots, leading to new insights into the unique characteristics of allosteric networks in various systems. However, DMS studies are mostly limited to the identification of phenotypes through the binary functional readout due to its high-throughput nature. Establishing the effect of a given mutation on protein function and other quantitative properties requires additional molecular details.

Another feature that is commonly observed in allosteric systems is the network of compensatory interactions,^42^ which leads to near-degenerate allosteric communication pathways. ^43^ This feature makes allostery highly adaptable during evolution and ensures robustness of functionally essential co-operativities;^37^ it can also facilitate re-wiring of allostery communication pathways in engineering applications.^12,44^ Nevertheless, these compensatory effects have major implications to the design of allosteric inhibitors, as the effect of design might be easily abolished by additional mutations.^4,6^ Therefore, robust designs should target allostery communications that, once disrupted, are difficult to restore. Such properties are costly to evaluate using experiments due to the need of sampling a large number of sequences. Providing insights into how easily perturbations due to allostery hotspot mutations are reversed using computational approaches^45^ is of tremendous value but has not been widely attempted. A fitting example in this context is a bacterial transcription repressor, tetracycline repressor (TetR). TetR’s normal function relies on the allosteric coupling between the ligand binding domains (LBDs) and the DNA binding domains (DBDs), which are separated by more than 20 Å (Fig. 1a). Binding of the inducer (ligand-Mg^2+^ complex) leads to the dissociation of the repressor from the promoter DNA sequence, allowing the access of RNA polymerase and therefore transcription of the repressed gene. Firmly establishing the contributions of hotspot residues has fundamental values to many aspects of prokaryotic physiology, including antibiotic resistance, and provides guidance to rational engineering of the TetR family for gene regulation.^46^ Therefore, allostery in TetR has been extensively studied with structural,^47–50^ mutagenesis^51–53^ and more recently, DMS studies.^37,38^ Based on propagation of structural distortions, a set of allostery hotspots abolishing the induction function in TetR were proposed^54^ and verified^55,56^ to localize at the domain interface of the two binding sites and relay the effect of inducer binding to the DBDs, whose reorientation impacts the DNA binding affinity. By sharp contrast, DMS analysis observed that allostery hotspots in the TetR family members are broadly distributed over a major portion of the protein structures,^37,38^ underscoring that a mechanical view of allostery,^23,57–59^ especially in TetR is incomplete. Moreover, the DMS experiments found that the loss of induction in hotspot mutants could be restored by additional mutations.^37^ These observations again highlighted that the allostery network has a considerable degree of degeneracy and redundancy, and not all allostery hotspots are of equal weights during evolution.

**Figure 1:**
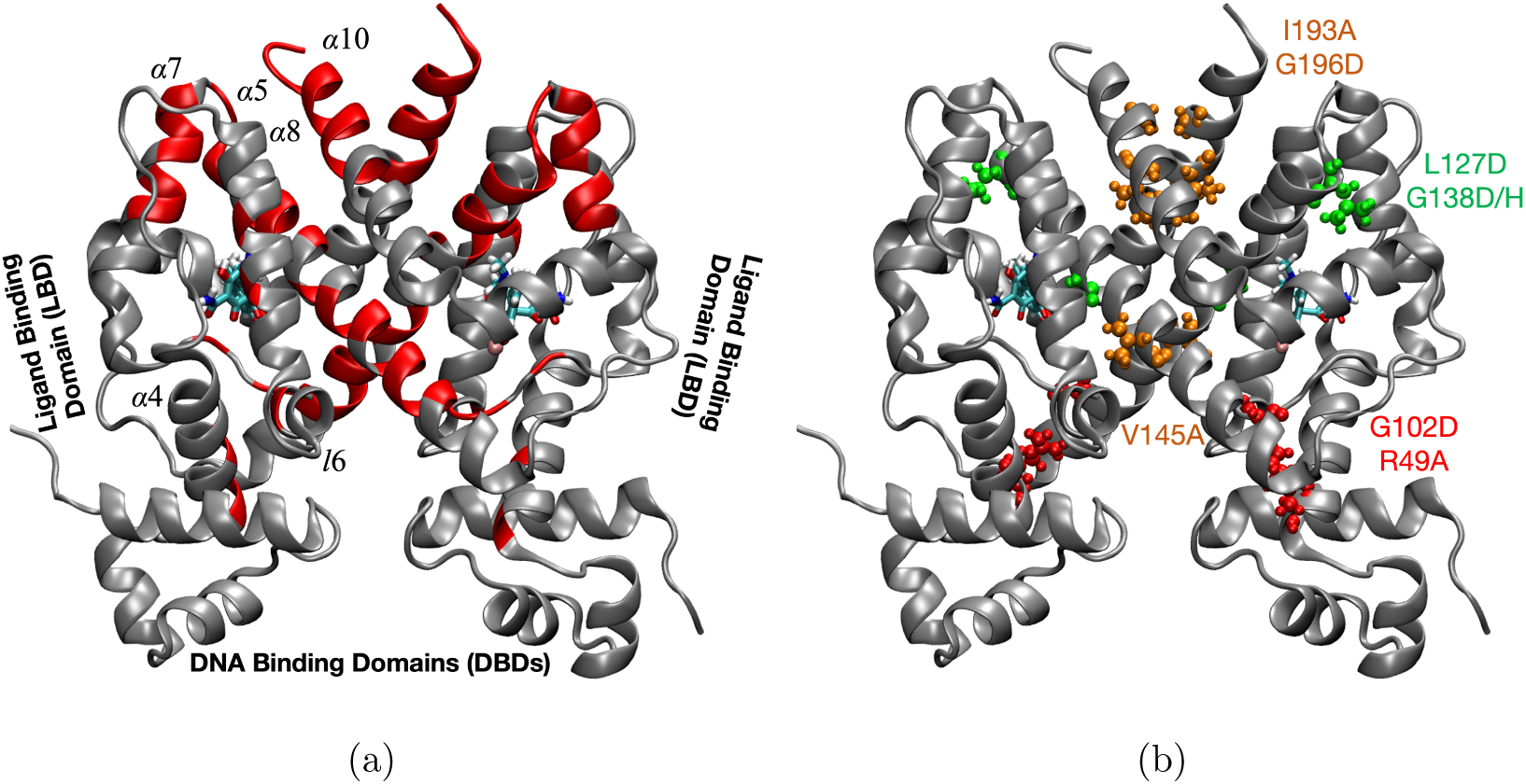
Overview of the TetR structure and experimentally-identified allostery hotspot distributions. (a) DMS studies^37^ revealed a broad distribution of allostery hotspots (in red) throughout the majority of the TetR structure. In the ligand-bound crystal structure (PDB code: 4ac0), the ligand, minocycline, is bound to a Mg^2+^ ion. The ligand is converted to anhydrotetracycline in all simulations. (b) The hotspot mutations studied in this work are grouped into three categories based on their locations: the domain interface (red), near the ligand binding site (green), and near the monomer-monomer interface (orange).

These considerations motivated our recent atomistic molecular dynamics (MD) simulations of the wild type (WT) TetR.^60^ Multiple local (e.g., backbone dihedral and residual contacts) and global (e.g., motional coupling and free energy landscapes) analyses established the molecular and energetic basis of the strong anti-cooperativity between the ligand and DNA binding sites through a conformational selection mechanism that underlies the allostery function of TetR. The multifaceted computational framework we established captured a major fraction of the hotspot residues identified in the DMS study and demonstrated that these residues contribute to allostery in distinct ways. Nevertheless, our previous simulations focused entirely on the WT protein, thus the roles and features of hotspot residues had to be inferred indirectly. Moreover, the general strategy for hotspot identification also generated a considerable number of “false positives” and missed hotspot residues in two regions, the middle of the *α*8 helix and the C-terminus of the protein.^60^ Evidently, the molecular mechanism of the disruption of allostery in the hotspot mutants, the non-inducible phenotype, remains unclear.

In the current study, the major goal is to establish a computational framework that enables us to evaluate the functional outcome of hotspot mutations in TetR. We show that such evaluation can be accomplished at qualitative (whether a mutant is non-inducible) and semi-quantitative (rank order the impact of the mutation) levels by judiciously combining extensive unbiased (6-60 *µ*s) and enhanced sampling molecular dynamics (MD) simulations. We select several representative mutants that involve hotspot residues in distinct regions of the protein (see Fig. 1b): domain interface (G102D, R49A), near the ligand binding site (L127D, G138D/H), and near the monomer-monomer interface, either in the middle of the *α*8 helix (V145A) or in the C-terminal helix (I193A and G196D) to explicitly probe their mutation effects.

## Computational Methods

### Unbiased MD Simulations of Different Functional States

Extensive unbiased MD simulations (6-60 *µ*s) are conducted to compare hotspot mutants and WT TetR in terms of key structural properties and construct free energy landscapes in multiple functional (apo, ligand-bound and DNA-bound) states on local NVIDIA P100 GPUs and the Anton 2 machine. WT trajectories were generated in the previous study.^60^ The mutants studied in this work include G102D, R49A, L127D, G138D, G138H, V145A, I193A, and G196D. All mutations are generated *in silico* based on the corresponding coordinates of the WT. Unbiased MD setup for the mutants, the definitions of various order parameters and structural properties are the same as our previous work on the WT.^60^ Simulation details and data analysis for the unbiased trajectories are presented in the **Supporting Information**. For the ligand-bound state, we start with the mutant ligand-bound structures based on the crystal structure of the WT (PDB code: 4AC0). For the DNA-bound state, the initial structure is modeled based on the crystal structure of the TetR(D)-DNA complex (PDB code: 1QPI), in which the protein has a high level of sequence homology to TetR(B) (66%). 1 *µ*s and 5 *µ*s trajectories are generated for the bound states on local GPUs and Anton 2, respectively.

For the apo state, three independent models are used as the initial structures to reduce bias and facilitate the exploration of the conformational space. The first structure is the same as the ligand-bound crystal structure but without the ligand-Mg^2+^ complex; the second structure is the same as the DNA-bound structure but without the DNA; the third structure is modeled using the TetR(D) apo crystal structure (PDB code: 1BJZ) as the template. Independent simulations of apo states are carried out both on local GPUs and Anton 2 starting from different initial structures for each system. For apo state simulations on local GPUs, 3 independent trajectories of 1 *µ*s or 3 *µ*s are carried out starting from the aforementioned three initial structures, respectively. For the apo state simulations on Anton 2, 3 independent trajectories of 15 *µ*s, 15 *µ*s, and 20 *µ*s are carried out for the WT starting from the three initial structures, respectively; 2 independent trajectories of 15 *µ*s each are carried out for G102D starting from the first two initial structures of the apo state, respectively; 3 independent trajectories of 10 *µ*s each are carried out for the rest mutants starting from the three initial structures, respectively.

### Metadynamics Simulations of Ligand-bound States

Multiple-walker well-tempered metadynamics simulations^61,62^ are carried out using PLUMED^63,64^ interfaced with OpenMM^65^ on NVIDIA A100 GPUs. The collective variable (CV), DNA-binding domain (DBD) distance, is the center-of-mass (COM) separation between the C*α* of residues 37-44 in each monomer. The bias factor is set to be 10. The simulation temperature is 303.15 K. A new Gaussian biasing potential is added every 1 ps with an initial height of 1.2 kJ/mol and a width of 0.2 nm. Ten walkers with different initial velocities are used per system in parallel while sharing hill history with other walkers every 1 ps. A harmonic restraint is applied to the CV with a force constant of 150 kJ/mol when the value of the CV is larger than 5 nm (upper wall) or smaller than 3 nm (lower wall). Other basic MD parameters are the same as the unbiased simulations. Convergence is monitored by the block-average analysis of which the details are presented in the next subsection. The simulation is stopped when adequate convergence has reached, which typically takes 110 - 140 ns per walker in each independent metadynamics run.

### Data Analysis for Metadynamics Simulations

Metadynamics trajectories are analyzed using the metadynamics reweighting algorithm ^66^ implemented in the PLUMED package (version 2.8)^64^ to obtain the unbiased distribution of the DBD distance, which is then converted into the 1D PMF. Block-average analysis is applied to assess convergence of metadynamics simulations and the associated statistical errors for each system. The average errors of the block-averaged PMF as a function of the block size are plotted as the convergence check. The final 1D PMF of each system is the block average of every 20 ns trajectories in each walker, and the shaded area in the PMF plot represents standard error of the mean (SEM) of the block-averaged PMF. The reweighted distributions of other order parameters, structural properties, as well as the 2D free energy landscapes are obtained using the same reweighting algorithm.

## Results

### Doubly-Bound States Are Better Stabilized in Hotspot Mutants

According to the induction function of TetR, ligand binding in the WT protein was proposed to rigidify the protein structure such that the two DBDs remain too far apart to allow favorable binding to the DNA. Thus, we reason that the loss of the induction function in some, if not all, hotspot mutants is due to adopting favorable conformations in the DBDs for DNA binding even in the ligand-bound state; i.e., these mutants are able to bind tightly to both the ligand and DNA in such conformations, referred to as the doubly-bound state. To test this hypothesis, we conduct metadynamics simulations to explicitly probe how hotspot mutations modulate the stability of the doubly-bound state, which is not expected to be stable in the WT protein. Under the condition of the DMS experiments, the ligand concentration is such that most TetR proteins are expected to be ligand bound. Therefore, free energy profiles for the DBD distance separation are computed in the ligand-bound state. The use of the DBD distance as the reaction coordinate is supported by previous structural studies of TetR^47,49^ and MD simulations on the order of 50-100 ns,^67^ which observed that the DBD distance can be used to distinguish induced (ligand-bound) and non-induced (DNA-bound) conformations, and thus a key indicator for the DNA binding affinity.

We note that the DBDs are perturbed by the hotspot mutations in both the DNA-bound (Fig. S1) and apo (Fig. S2) states in unbiased simulations. As shown in Fig. S1, the DNA-bound state of the mutants tends to span broader DBD distances compared to the WT, leading to Kullback-Leibler (KL) divergence values in the range of 0.8-1.8 for the corresponding DBD distance distributions (Table S2). These values reflect modest differences in the DBD conformations between the mutants and WT. Regarding the DBDs in the apo state, their conformations in the hotspot mutants are overall similar to the WT, except that the DBD separations are slightly shifted towards longer distances (Fig. S2), most notably in G138D and G138H.

In the WT, the computed free energy profile (Fig. 2) indicates that there is a single free energy basin at *∼* 46 Å along the DBD distance coordinate. The basin (ligand-only basin) corresponds to the crystal structure of the ligand-bound TetR, in which the DBD distance is *∼* 42 Å.^47^ As the DBD distance decreases, the free energy increases monotonically. For the DBD distance reaching *∼*38 Å, the corresponding conformation is expected for favorable DNA binding,^49^ which has a free energy cost about 10 *k_B_T* . These observations are consistent with the normal function of TetR. The low DNA binding affinity upon ligand binding suggests that compressing the DBDs in the ligand-bound state is energetically unfavorable in the WT. In all the hotspot mutants studied, by remarkable contrast, the computed free energy profiles are double-well in nature (Fig. 2a-c). The ligand-only basin is consistently observed at the large DBD distance of *∼*46 Å, similar to the WT. In addition, a second free energy basin is observed at a shorter DBD distance expected for favorable DNA binding in the ligand-bound state. The location of this inner basin is largely consistent (*∼*38 Å) among all the mutants studied here.

**Figure 2:**
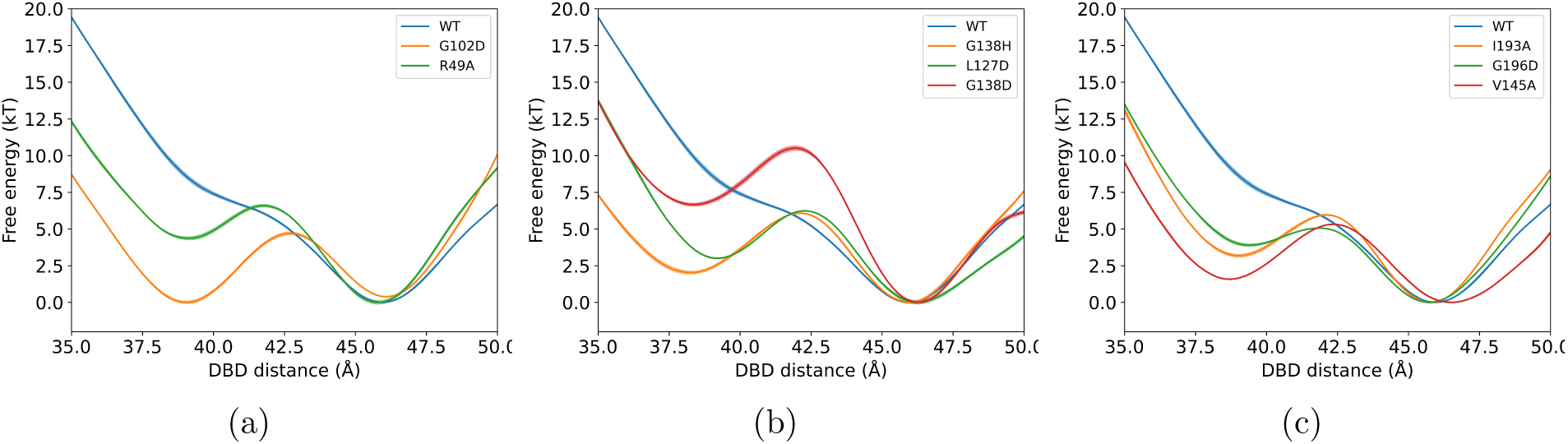
Potentials of mean force (PMF) for the DBD distance separation in the ligand-bound states from metadynamics simulations. The reaction coordinate is the center of mass separation between the two DBDs in each monomer. See the precise definition in Materials and Methods. All PMFs show a basin at *∼* 46 Å, referred to as the “ligand-only basin”. All mutant PMFs also show another basin at *∼* 38 Å, referred to as the “doubly-bound basin”. Subfigures are grouped by the regions shown in Fig. 1b. The solid lines are the average values and the shaded areas show the statistical error estimated from block analysis. The statistical errors are all on the order of 0.1 *k_B_T* . For additional analysis of statistical error and convergence, see Fig. S3 in the **Supporting Information**.

To further verify the nature of the second basin in the PMFs along the DBD distance in the mutants, we measure the KL divergence of the DBD distance distributions between the ligand-bound and DNA-bound state simulations (see Figs. S4-S5). As shown in Table 1, the WT simulations lead to larger KL divergence values of the DBD distance distributions between the ligand-bound and DNA-bound states. A smaller KL divergence value indicates a larger similarity between the two distributions under comparison, suggesting that the DBD distance distributions in the ligand-bound state of the mutants are closer to the corresponding distributions in the DNA-bound state such that the mutants are more favorable for DNA binding than the WT in the ligand-bound state. It is also worthwhile decoupling the contributions of the two basins from metadynamics simulations to better illustrate the origin of more favorable DNA binding in the ligand-bound state of the mutants. Accordingly, we further split the reweighted distributions from metadynamics simulations into two components at the DBD distance of 42.5 Å as the boundary. The DBD distance distributions of the inner basin in the ligand-bound state of the hotspot mutants overlap well with the corresponding DNA-bound state with KL divergence values no larger than 1.0, showing that the DBDs in the inner basin indeed favor DNA-binding. The large KL divergence values in the basin at the longer DBD distance suggest that the DNA is unfavorable to bind, verifying the nature of this ligand-only basin. Another structural variable thought to be important to DNA binding (affinity) is the relative angle of the *α*4 helices at the domain interface connecting the DBDs and LBDs^47,49^ (see definitions of key DBD order parameters in SI). KL divergence of the *α*4 angle distributions in the two bound states is also substantially lower in the hotspot mutants than in the WT (Table S3 and Fig. S6-S7), also showing that the ligand-bound states of the mutants have higher populations in the DNA-binding-competent conformation than the ligand-bound state of the WT. Collectively, these results quantitatively demonstrate that the free energy basin at the short DBD distance sampled in the metadynamics is indeed favorable for DNA binding in the ligand-bound state. Therefore, although only the ligand is explicitly included in the metadynamics simulations, we refer to this basin as the “doubly-bound basin” since the corresponding conformational ensemble is expected to bind tightly to both the ligand and DNA.

**Table 1:** Overlap of DBD distance distributions between the ligand-bound and DNA-bound states characterized by the Kullback-Leibler (KL) divergence *^a^*

|  | WT | G102D | R49A | G138H | L127D |
| --- | --- | --- | --- | --- | --- |
| reweighted | 6.8 | 1.2 | 3.0 | 1.7 | 2.8 |
| doubly-bound basin<sup>b</sup> | 6.0 | 0.4 | 0.5 | 0.1 | 0.6 |
| ligand-only basin<sup>b</sup> | 54.4 | 56.7 | 57.3 | 56.7 | 57.5 |
|  | G138D | V145A | I193A | G196D |  |
| reweighted | 3.8 | 2.8 | 2.0 | 3.3 |  |
| doubly-bound basin<sup>b</sup> | 0.4 | 1.0 | 0.8 | 1.0 |  |
| ligand-only basin<sup>b</sup> | 56.7 | 57.0 | 56.1 | 57.2 |  |

b. Reweighted distributions from metadynamics simulations are split into “doubly-bound basin” and “ligand-only basin” components at the DBD distance of 42.5 Å as the boundary.

Among the hotspot mutants studied, the doubly-bound basin is higher in free energy than the ligand-only basin at the longer DBD distance by 2-7 *k_B_T* . The only exception is G102D, in which the two basins are similar in stability (Fig. 2a), suggesting that G102D is highly poised to bind both the ligand and the DNA. While the R49A mutation is also close to the domain interface and involves a similar magnitude change in net charge, the mutation effect on the DBD energetics is notably smaller. Interestingly, G138D and G138H exhibit rather different stabilities of the doubly-bound basins (Fig. 2b) despite implicating mutation of the same residue. These variations underscore the importance of the unique local interactions associated with specific mutations. For example, in G102D, the introduced aspartate D102 is engaged in new interactions (compared to the WT) with several cationic residues nearby, such as K155’ and R158’ in *α*8’, and R49 in *α*4 (Fig. S8). By contrast, in R49A, the interaction between R49 and E157’ in *α*8’ in the WT^60^ is lost. As shown more explicitly in Fig. S9, the top ten contact changes in G102D are localized to the domain interface, implicating mainly the end of the *α*8 helix and the top of the *α*4 helix, along with loop *l*6. In R49A, while the top ten contact changes also involve residues in the loop *l*6 and top of the *α*4 helix, the changes are more delocalized into the lower regions of the DBDs. In G138D, the introduced aspartate D138 is engaged in hydrogen-bonding interactions with the hydroxyl group in the ligand and therefore has rather static orientations. While in G138H, the lack of such hydrogen-bonding interactions leaves the introduced histidine H138 flexible, with occasional flip of the sidechain to push on *α*5 (Fig. S8). Finally, it is worth noting that while previous WT simulations did not identify any distinct properties of the hotspot residues in the middle of *α*8 and C-terminal region (*α*10),^60^ the current metadynamics simulations of the *α*8 mutant (V145A) and C-terminal mutants (I193A and G196D) have clearly captured the effect of the mutations on the DBDs (Fig. 2c). In these three mutants, the doubly-bound basin is distinctly visible and is only higher in free energy by *∼*2-4 *k_B_T* than the ligand-only basin.

### Inter-Domain Coupling Is Reduced in Hotspot Mutants

The normal induction function of the WT TetR relies on the strong anti-cooperativity between the LBDs and DBDs. The double-well feature of the computed DBD free energy profiles of the hotspot mutants in Fig. 2 suggests that the magnitude of inter-domain coupling is substantially reduced compared to the WT. As a result, these mutants are more inclined to bind tightly to DNA even in the ligand-bound state. Importantly, the reduction of inter-domain coupling is also evident from several additional lines of analyses, further supporting the robustness of the conclusion.

First, we note that, in the ligand-bound state simulations, the ligand binding pocket in the hotspot mutants exhibits notable changes depending on the DBD distance. At a longer DBD distance in the ligand-only basin, the l6-*α*8 distance (the C*α* distance between R104 and L134) that characterizes the compactness of the binding pocket exhibits overall similar distributions as the WT (Fig. 3a-c), with merely shifted peak positions or modified distribution widths. In the doubly-bound basin, which features a compressed DBD distance, much larger changes in the ligand binding pocket distributions relative to the WT are observed for several mutants (Fig. 3d-f). For example, both G102D and G138H exhibit much broader distributions. The peak position is also notably shifted in L127D, the two G138 mutants G138H/D, as well as V145A and G196D (also see Fig. S10 and Table S4). These observations suggest that the coupling between the binding pocket and the DBDs is not altered much in the ligand-only basin in the hotspot mutants, but the coupling is more significantly modified in the doubly-bound basin of the mutants. In particular, the substantially expanded pairwise distance distributions in G102D and G138H in the doubly-bound basin suggest that the coupling between LBDs and DBDs is much reduced relative to the WT. The reduced inter-domain couplings at compressed DBD distance are consistent with the higher stability of the doubly-bound basin in these hotspot mutants in comparison to the WT (Fig. 2).

**Figure 3:**
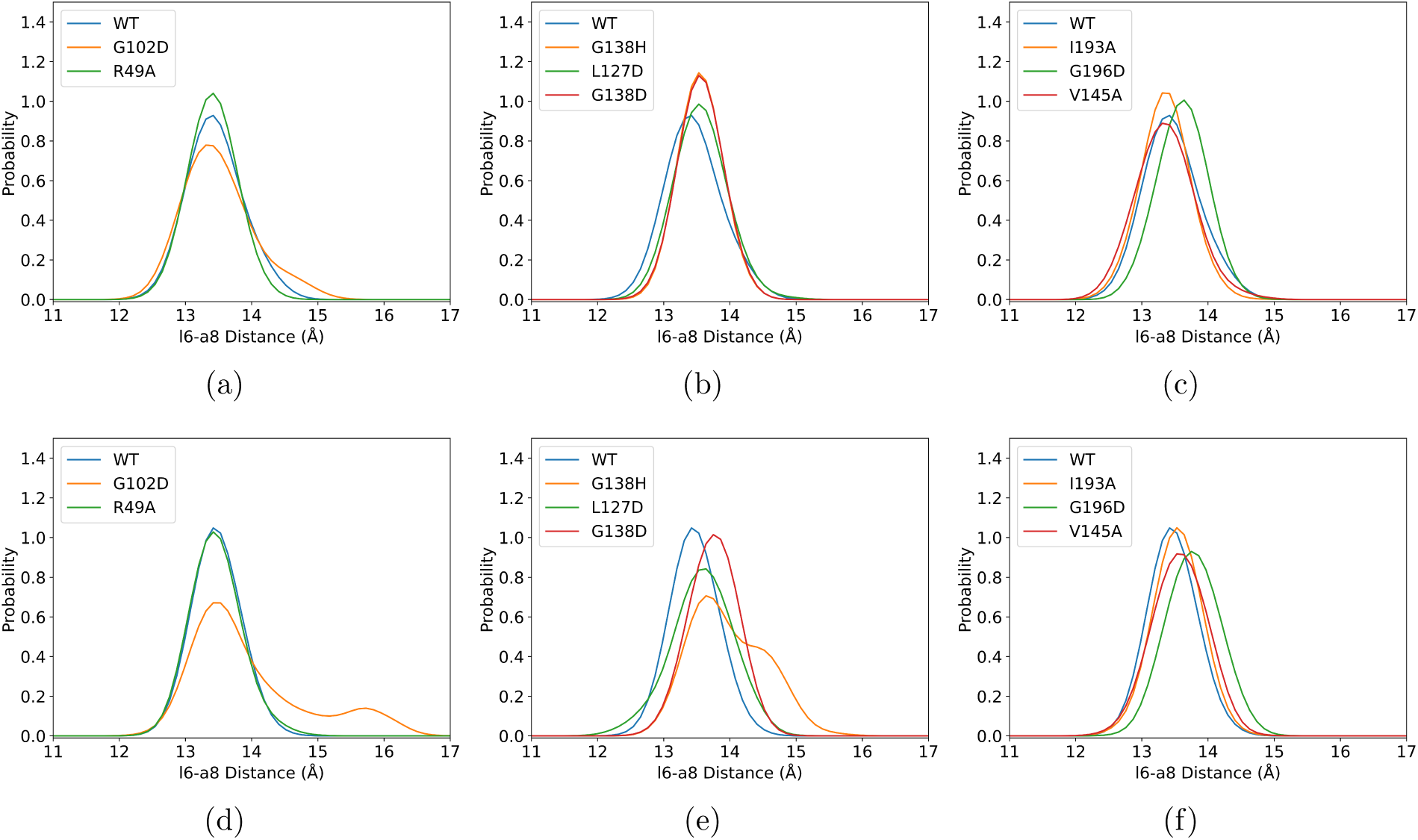
Binding pocket distance distributions in different conformational states of the ligand-bound states from metadynamics simulations. Subfigures are grouped by the regions shown in Fig. 1b. The reweighted distributions of the C*α* distance between R104 and L134 (l6-*α*8 distance. l6 stands for the linker region of *α*6-*α*7.) (a-c) in the “ligand-only basin” (DBD distance range is between 46.5 Å and 47 Å), and (d-f) in the “doubly-bound basin” (DBD distance range is between 37.5 Å and 38 Å) from metadynamics simulations.

The variation of inter-domain coupling among the hotspot mutants relative to the WT is also directly visualized in the free energy landscapes of the ligand-bound states projected onto the two-dimensional space spanned by the DBD distance and the l6-*α*8 distance (Fig. 4). In the WT, free energy landscapes of the two bound states are distinct and exhibit limited overlap along either collective variable in the unbiased simulations as well as in the reweighted landscapes from metadynamics simulations, reflecting the strong anti-cooperativity between the LBDs and DBDs. In the free energy landscapes of the mutants, especially in the ligand-bound simulations, the DBD distance distributions are much broader than the WT. Two basins are clearly observed along the DBD distance coordinate in the reweighted landscapes from metadynamics simulations, while they share very similar equilibrium l6-*α*8 distances. These observations reflect weaker inter-domain anti-cooperativity in the mutants. Furthermore, along the l6-*α*8 distance coordinate for G102D and G138H (Fig. 4b, d), the binding pocket distributions are much broader with substantial degrees of overlapping at the compressed DBD distance, suggesting that the ligand binding pocket becomes even more flexible when the DBDs favor DNA binding and thus showing even weaker inter-domain coupling in the doubly-bound basin in these mutants.

**Figure 4:**
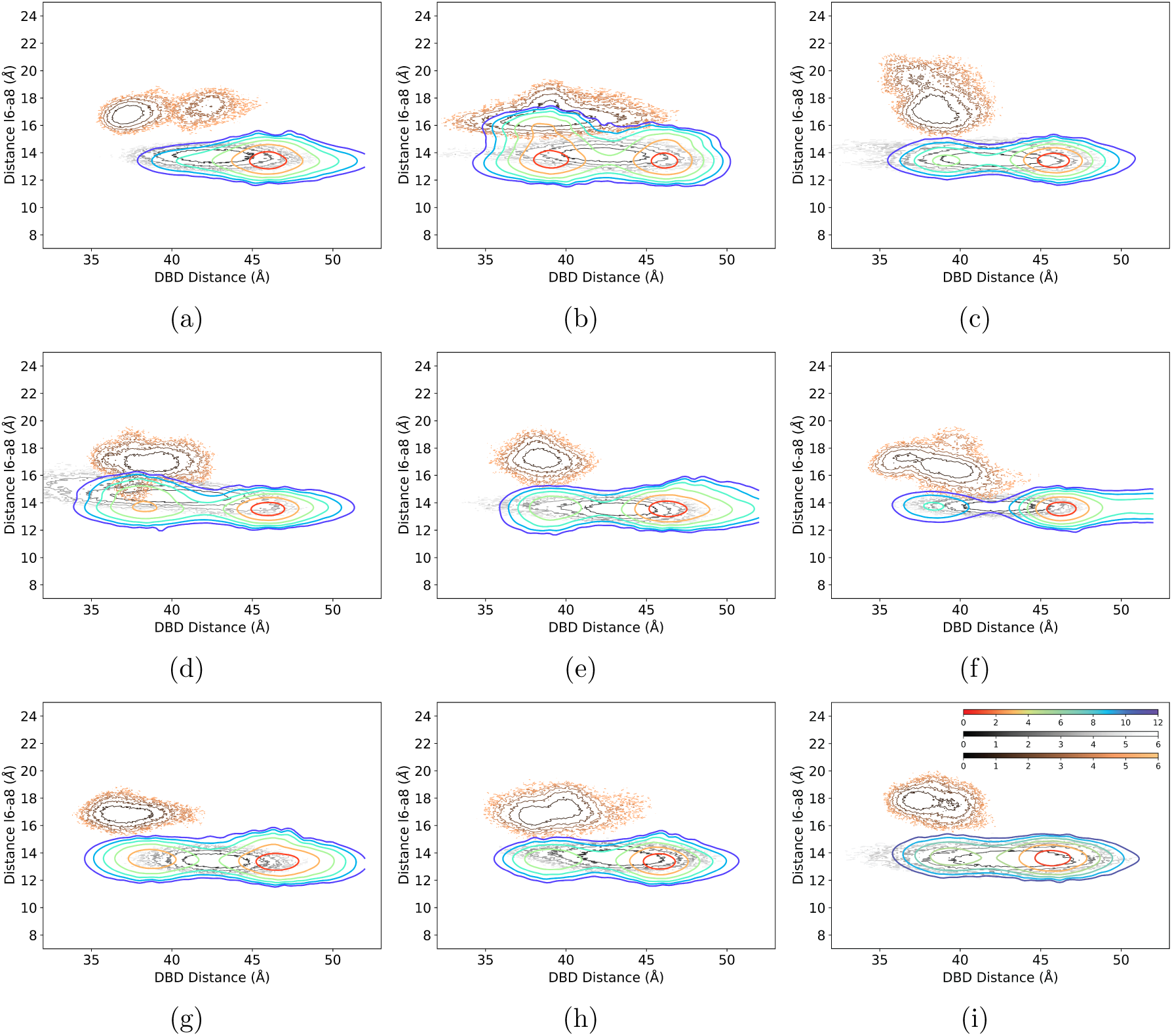
Two-dimensional free energy landscapes of the DNA-bound and ligand-bound states: (a)WT, (b) G102D, (c) R49A, (d) G138H, (e) L127D, (f) G138D, (g) V145A, (h) I193A, (i) G196D. The horizontal axis is the center-of-mass distance between the DBDs, and the vertical axis is the C*α* distance between R104 and L134 (l6-*α*8 distance), which characterizes the compactness of the ligand binding pocket. For the DNA-bound states, the results are based on unbiased MD simulations (see Table S1 for simulation length summary). For the ligand-bound states, results based on unbiased MD simulations and reweighted distribution from metadynamics simulations are shown for comparison. While the unbiased and metadynamics simulations sample mostly similar conformational ensembles, due to the free energy barrier along the DBD distance (Fig. 2), the metadynamics simulations better capture the underlying free energy landscape and structural properties, especially for the “doubly-bound basin”, which is hardly sampled in the unbiased simulations at the microsecond time scale. Color scheme: rainbow - ligand-bound state metadynamics simulations; gray - unbiased ligand-bound state simulations; copper - DNA-bound simulations. Color bars are in the unit of *k_B_T* .

Next, we examine the free energy landscapes of the apo states projected onto the two-dimensional space spanned by the collective DBD motion (represented by the first principle component PC1 used in our previous work^60^), and *α*7*−α*8*^′^* distance (the C*α* distance between Q109 and E147’) of the ligand binding pocket (Fig. 5a-c). Analysis of the apo landscape is complementary to the bound-state simulations discussed so far, as the shapes of the apo landscapes reflect the intrinsic features of inter-domain coupling in the absence of any local stabilization by the binding event of the ligand or the DNA. The two-dimensional free energy landscape of the WT shown in Fig. 5a is much broader than those computed of the bound states,^60^ pointing to a conformational selection mechanism for TetR function. Moreover, the diagonal shape of the landscape is consistent with the strong anti-cooperativity expected for the LBDs and DBDs. For the studied hotspot mutants, the apo free energy landscapes are similarly broad, indicating that the conformational selection mechanism is still applicable. The diagonal shapes of the landscapes remain qualitatively similar to the WT for most mutants (Fig. S11), suggesting that the anti-cooperativity between the LBDs and DBDs is not significantly perturbed in the apo state of these mutants.

**Figure 5:**
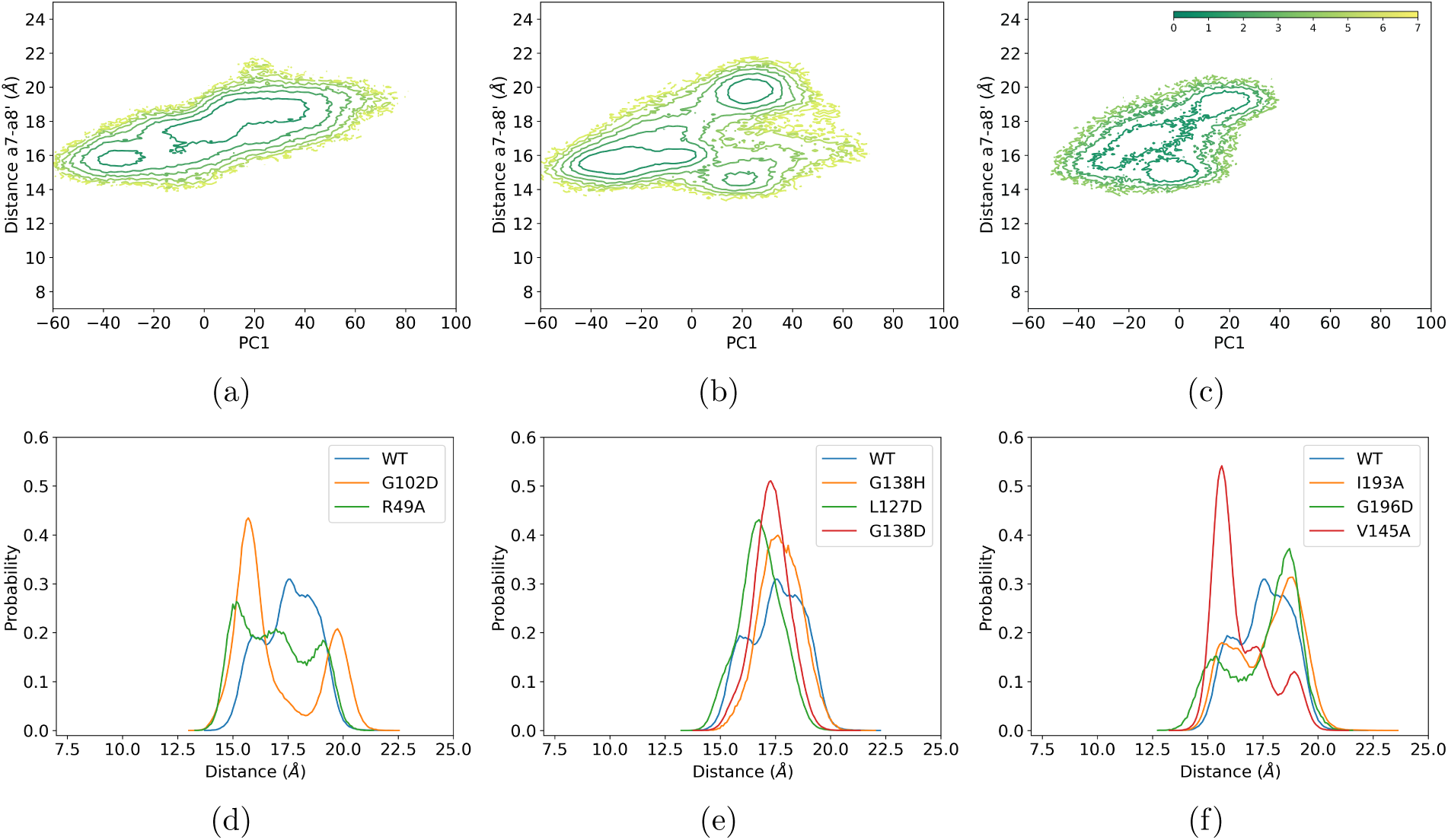
Free energy landscapes and ligand binding pocket distance distributions of the apo states. Two-dimensional free energy landscapes of the (a) WT, (b) G102D and (c) R49A variants. Results of the remaining mutants are qualitatively similar to the WT and shown in Fig. S11. The color bar is in the unit of *k_B_T* . The horizontal axis is the first principal component (PC1), and the vertical axis is the C*α* distance between Q109 and E147’ (*α*7-*α*8’ distance). PC1 describes the complex motions of the DBDs, while the inter-helical distance characterizes the compactness of the ligand binding pocket. The WT and mutants generally feature diagonal shape landscapes, while an extra basin is also observed in the mutants (G102D and R49A). Two-dimensional free energy landscapes projected onto different order parameters are shown in Fig. S12, which also exhibit similar changes for the two domain-interface mutants (G102D and R49A). (d-f)Distributions of the *α*7-*α*8’ distance in different mutants. The results are based on extensive unbiased samplings (9-60 *µs*) for each variant. Subfigures are grouped by the regions shown in Fig.1b.

Exceptions are the two mutations at the domain interface, G102D and R49A, of which the free energy landscapes deviate significantly from the diagonal shape with an additional basin that features a positive PC1 value and a short *α*7 *− α*8*^′^* distance (compare Figs. 5b,c and 5a), which corresponds to a conformational state that has low binding propensity to both the ligand and DNA. Importantly, the most evident changes in the free energy landscapes associated with decreased inter-domain coupling in G102D and R49A are also observed when the free energy landscapes are projected onto other two-dimensional spaces spanned by the DBD distance and the *α*7 *− α*8*^′^* distance (Fig. S12). As explained in the **Supporting Information** using a revised thermodynamic model (Fig. S13) for allostery in TetR,^60^ such changes in the free energy landscape are strong signatures for the decrease of inter-domain coupling. The apo landscape in the WT features two basins that correspond to the induced (ligand-bound, *L_B_* · *D_E_*) and non-induced (DNA-bound, *L_E_* · *D_B_*) conformations. The inter-domain coupling free energy, *γ* explicitly controls the anti-cooperativity which disfavors the two domains simultaneously adopting either the high-binding-affinity (*L_B_*· *D_B_*) or the low-binding-affinity (*L_E_* · *D_E_*) conformations in the WT. With reduced *γ* by mutations (Fig. S13c), a third basin representing the *L_E_* · *D_E_*conformation with a lower free energy is observed in the apo landscapes of G102D and R49A. Note that the *L_B_* · *D_B_* conformation still remains higher in free energy in the apo state, although the doubly-bound state is further stabilized by the binding of ligand and DNA and therefore expected to become the dominant population for the corresponding mutants in the presence of ligand and DNA (Fig. S13c). Indeed, the doubly-bound conformation is clearly visible as a free energy basin in the computed landscapes for the ligand-bound state of hotspot mutants (Fig. 4b-i).

### Binding Pocket Is Perturbed to Decrease Ligand Binding in Hotspot Mutants

The free energy results (Fig. 2) and analysis of inter-domain coupling suggest that the studied hotspot mutants have much stronger propensities to bind both the ligand and DNA, which is consistent with their non-inducible phenotype observed in the DMS experiment.^37^ However, a higher population of the doubly-bound state is not the only potential cause for the loss of induction function. Reducing TetR’s apparent ligand binding affinity to be lower than the binding affinity for the DNA promoter sequence, or enhancing the DNA binding affinity would also lead to the non-inducible phenotype (see Fig. S13). In this context, we note that the apparent binding affinity in general contains two contributions, the free energy cost of transitioning from the inactive (binding-incompetent) to the active (binding-competent) conformation, and the intrinsic binding affinity of the active conformation. Moreover, the apparent ligand or DNA binding affinity can be modified by hotspot mutations through perturbations in either the ligand/DNA-bound state or the apo state, or both. Perturbation of the bound conformational ensemble modifies the intrinsic binding affinity. For the apo state, changes in the active conformation also modify the intrinsic binding affinity, while population shifts in the equilibrium between the inactive and active conformations lead to variations in the activation free energy component of the apparent binding affinity. Therefore, we evaluate the impact of the hotspot mutations on the properties of both bound states and the apo state.

For the DNA-bound states, the ligand binding pocket remains loosely packed in both WT and the hotspot mutants (Fig. S14-S16). For the ligand-bound states, at long DBD distances (in the ligand-only basin), the ligand binding pocket remains tightly packed and also experiences limited perturbation by mutations (Fig. 3a-c). With compressed DBD distances (in the doubly-bound basin), however, considerably broader binding pocket distributions are observed in several mutants (Fig. 3d-f), especially in G102D and G138H. Therefore, upon DNA binding, the intrinsic binding affinity to the ligand is expected to be further weakened in these mutants as compared to the WT, which contributes to the diminished induction activities.

In the apo states, many hotspot mutations impact the conformational ensembles of the ligand binding pocket. As shown in Fig. 5d, the WT ligand binding pocket favors the binding-competent conformation (long *α*7-*α*8’ distances) in the apo state. In the two mutants at the domain interface (G102D and R49A), by contrast, the ligand binding pocket is dominated by the binding-incompetent conformation (short *α*7-*α*8’ distances) that is favored in the DNA-bound state (see Fig. S14); a similar trend is also observed for V145A (Fig. 5f), in which the mutation is close to both the domain interface and the monomer-monomer interface (see Fig. 1b). Therefore, ligand binding in these mutants requires a notable free energy cost for the binding pocket to adopt the binding-competent conformation, leading to a lower apparent ligand binding affinity relative to the WT. In several cases, such as G102D and C-terminal mutants (I193A and G196D), the peak positions of the distributions favoring ligand-binding are substantially shifted (Fig. 5f), suggesting notable change in the binding-competent conformation and therefore variation in the intrinsic binding affinity. In other cases, such as L127D (Fig. 5e), the *α*7-*α*8’ distance distribution becomes unimodal and the peak position shifts towards a short distance that is favored in the DNA-bound state (also see Fig. S14), also suggesting considerable conformational rearrangements in the binding pocket and reduction in the ligand binding affinity.

## Discussion

### Evaluation of Functional Outcome of Hotspot Mutations in TetR

Evaluation of hotspot mutations is of major interest to applications that aim to modulate allostery because hotspot residues dictate the degree of allosteric coupling and therefore function of allostery systems. In our recent study,^60^ motivated by the broad distribution and distinct contributions from allostery hotspots, we emphasized the importance of analyzing multiple properties at both the local and global scales, which was demonstrated to be essential to identifying multiple hotspots. However, since only the WT protein was investigated, contributions of residues to allostery can only be inferred indirectly from WT simulations. The overarching objective of this study, which has been rarely demonstrated in previous computational studies of allosteric system, is to establish a computational framework to evaluate the functional outcome of hotspot mutations in TetR, both qualitatively (whether a mutant is non-inducible) and semi-quantitatively (rank order the impact of the mutation and infer the degree of rescuability). We combined extensive (up to 60 *µs*) atomistic MD simulations, systematic analysis of structural and dynamical properties and multiple modest-length (*∼ µs*) enhanced sampling simulations to study WT and several representative mutants in multiple functional states.

The current work demonstrates that functionally relevant PMF simulations are effective at capturing mutation effects in various hotspot mutants. We first identify the DBD distance as a collective variable that can be used to distinguish inducible and non-inducible TetR conformations, which is supported by previous structural and simulation studies. Accordingly, free energy profiles along this collective variable are highly correlated with the DNA binding affinity and the induction function of TetR. Encouragingly, all mutant simulations convincingly captured the mutation effects, including those missed in previous WT simulations. A second basin at a short DBD distance is only consistently observed in the hotspot mutants. KL divergence analysis clearly indicated that this second basin corresponds to a conformation that favors DNA binding in the ligand-bound state; i.e., the conformation is capable of binding to both ligand and DNA (the doubly-bound state). Stabilization of the doubly-bound state in these hotspot mutants explains their loss of induction function. Moreover, the free energy simulations revealed different degrees of stabilization of the doubly-bound state and underscored the importance of the unique local interactions associated with specific mutations. This further highlights the value of mechanistic analysis of allostery, which facilitates the identification of proper coordinate(s) for functional free energy landscape computations. As alternative approaches that rely minimally on prior knowledge of the system, creative computational methodologies based on machine learning analysis of conformational ensembles associated with distinct functional states^68–71^ represent promising directions to pursue.

The successful recapitulation of the hotspot mutation effects underscores the value of conducting explicit mutant simulations for the evaluation of hotspot candidates, since mutant simulations are able to capture the cascade of changes in structure and energetics not evident in the WT. Additionally, compensation effects of nearby residues are included, and therefore “false positives” can be minimized. In terms of the scope of the hotspot mutants, we have explored numerous mutations that are located in distinct structural regions of the protein and shown consistent trends in the shift of the free energy profile, further supporting the relevance of such computations to the evaluation of functional outcome. As complements to enhanced sampling, extensive unbiased bound state simulations are important for providing a valid reference for the analysis of the nature of free energy basins emerged in enhanced sampling simulations (e.g., see Table 1). Apo state MD simulations also reflect the intrinsic energetic coupling between the ligand and DNA binding domains to demonstrate the conformational selection mechanism of allostery in both the WT and mutants.

### Rescuability of Allostery Hotspots

In the DMS study of TetR,^37^ another remarkable observation was that the loss of induction function in hotspot mutants could be rescued by additional mutation(s), also in a distributed and degenerate fashion. At the qualitative level, since the induction function can be disrupted in multiple ways by perturbing either inter-domain coupling or intra-domain energetics,^60^ we expect that restoration of the normal function can also be accomplished in degenerate fashions through reversing changes in these properties. At the quantitative level, the degree of rescuability was defined as the ability of restoring the TetR function on non-inducible (dead) mutants. The DMS experiments found that among the five hotspots analyzed experimentally, the degree of rescuability follows the order of N129D*>*G196D*>*D53V*>*R49A*>*G102D.^37^ As discussed in our earlier work,^60^ successful design of allostery inhibition^2,4^ needs to target allostery hotspots that are particularly difficult to rescue, since otherwise allostery can be readily restored with additional mutations, leading to abolishment of inhibition. Therefore, being able to evaluate the rescuability of allostery hotspots using computation is of great value.

In this context, a closely related quantity is the magnitude of perturbation on the free energy landscape in the ligand-bound state with their degree of rescuability. The G102D mutant, in which the doubly-bound basin is similar in stability to the ligand-only basin; G102D was also observed to be most difficult to rescue in the DMS analysis. Although R49A is also a mutation at the domain interface and features an identical change in the net charge, the magnitude of the effect on DBD energetics is notably smaller, which is consistent with the higher degree of rescuability of R49A observed in the DMS experiment.^37^ As discussed above, the impact of mutation can be substantially different due to the specific local interactions unique to each mutation site (see Fig. S8-S9). Moreover, the relative stability of the doubly-bound basin and the ligand-only basin is comparable in G196D and R49A. This is qualitatively consistent with the similar degrees of rescuability of these two mutants observed in the DMS experiment. Therefore, these observations support that mutations that lead to a larger change in the DBD free energy landscapes are more difficult to rescue.

The general consistency between computational and experimental results regarding mutant rescuability suggests that it is feasible and meaningful to use functional free energy simulations to rank hotspot mutations for inferring rescuability. Indeed, the considerable level of redundancy for the allostery network identified in DMS studies raised the questions of whether engineering allosteric modulators can be a successful path. The current work helps compare the rescuability of hotspot mutations and supports that engineering designs to modulate allostery could be tested through MD simulations. Our computational strategies take advantage of multiple, modest-length enhanced sampling simulations and therefore are scalable to other allostery systems. Indeed, on modern computational hardware, multi-walker metadynamics simulations using well-defined collective variables converge rather efficiently on the microsecond time scale (see, for example, convergence analysis in Fig. S3) and therefore are readily affordable for screening a considerable number of mutants for functional evaluations.

Another popular and computationally efficient strategy for dissecting protein allostery involves analyses that target communication pathways such as suboptimal path calculations.^17–22^ As presented and discussed in the **Supporting Information**, while they can reveal differences between mutants and WT proteins, it is difficult to draw any firm conclusion regarding the functional impact of mutations. For example, we do not observe any clear correlation between the path-length distribution (Fig. S18) or the number of high-occurrence residues (Fig. S20) with the induction function of the TetR proteins studied here. Thus, the additional computational cost associated with functional free energy simulations is fully justified for the evaluation of allostery modulations.

### The Mechanism of Allostery Modulation by Hotspot Mutations

The extensive analyses of structural features of different functional states of the mutants and 2D free energy landscapes provided further insights into the molecular mechanisms through which the induction function is perturbed by the hotspot mutations. We illustrate that the loss of induction function in the hotspot mutants is due to either reduced inter-domain coupling, which leads to stabilization of the doubly-bound state, or perturbation of the ligand binding pocket conformation, which reduces apparent ligand binding affinity. A single mutation may lead to notable perturbations in both types of properties, such as in G102D. Therefore, the same non-inducible phenotype could be the result of perturbations on distinct structural and energetic properties. Our analysis of multiple functional states or free energy basins further reveals that the inter-domain coupling is not simply a constant value but dependent on the particular conformational and binding state of the protein. The apparent binding affinity can also be modulated by perturbing either the intrinsic binding affinity associated with the binding-competent conformations, or the activation free energy component, which depends solely on the conformational distribution of the binding site in the apo state.

Therefore, these observations generally support the thermodynamic model proposed in our previous work.^60^ It classifies allostery hotspots into two distinct types that dictate inter-domain coupling and intra-domain energetics, respectively. The current analysis of the mutants at the atomic level highlights that a single mutation may lead to notable perturbation in both types of properties; for example, G102D clearly impacts both inter-domain coupling and conformational distributions of the ligand-binding pocket. Therefore, the type 1 & 2 hotspots discussed in the context of the thermodynamic model^60^ are *idealized* classifications and the realistic situations tend to mix both effects. The PMF results for the DBD separation in the ligand-bound states (Fig. 2) clearly demonstrate that hotspot mutation effects can propagate from many regions in the protein to the DBDs, including the C-terminus, which is at the opposite end of the protein structure to the DBDs. This observation highlights the global nature of energetic coupling in proteins,^72^ and is consistent with the distributed nature of allosteric networks as observed in the DMS study^37^ and our previous MD analysis.^60^ These general features are broadly applicable to many allosteric systems, especially those sharing the two-domain structural topology. To further quantify the relative contributions of inter-domain and intra-domain effects, additional analysis of experimental data such as the induction activity as a function of the ligand concentration is required.^73^ Moreover, our mechanistic analysis relies on equilibrium computations and therefore does not aim to probe any causal relationship among different perturbations, the exploration of which requires non-equilibrium simulations.^74^

The essence of our dissection of allostery modulation in TetR concerns the analysis of inter-domain coupling and intra-domain properties that affect the apparent binding affinity of effector (inducer) on an equal footing. The magnitude of inter-domain coupling explicitly dictates the degree of anti-cooperativity between ligand and DNA binding, while the apparent effector binding affinity also influences the competition between ligand-bound and DNA-bound populations and therefore the degree of inducibility of TetR. Since the proper function of most allostery systems relies on co-operativity between distinct domains and tight binding of the effector, the strategies and insights we have established through the analysis of TetR have major implications to the general prediction and evaluation of allostery hotspots in proteins. This realization offers insights into how distal mutations might affect inhibitor potency in the context of drug resistance development.^6^ Moreover, it also highlights additional flexibility in the strategies that can be used to fine-tune the various properties relevant to the allostery function, including effector specificity.^44^

## Conclusions

For many biomedical and biotechnological applications, modulating allosteric coupling offers unique opportunities. Such efforts can largely benefit from the efficient prediction and evaluation of allostery hotspot residues that dictate the degree of co-operativity between distant sites. In several hotspot mutants revealed by recent deep mutational scanning (DMS) experiments of a bacterial transcription factor, tetracycline repressor (TetR), we demonstrate that effects of allostery hotspot mutations can be evaluated by judiciously combining extensive unbiased and enhanced sampling molecular dynamics simulations. The results recapitulate the qualitative effects of these mutations on abolishing the induction function of TetR and provide a semi-quantitative rationale for the different degrees of rescuability to restore allosteric coupling of the hotspot mutations observed in the DMS analysis.

Free energy landscapes and variations of structural and energetic properties of different functional states of the mutants show that the hotspot mutations perturb inter-domain coupling and/or intra-domain conformational properties relative to the wild type. Reduced anti-cooperativity between the distal binding sites is found in the mutants to reflect the loss of function. Multiple mutations also alter the conformational ensemble of the ligand binding site for weakened binding, suggesting that hotspot mutations can be used to fine-tune allostery properties including effector specificity. Several mutations clearly perturb multiple properties, also highlighting the complexity of realistic situations. Thus, the same non-inducible phenotype could be the result of perturbations in distinct structural and energetic properties. Overall, The results support our mechanistic model that hotspot residues contribute to allostery through distinct molecular mechanisms, a feature that we propose to be broadly applicable to many allosteric systems. For effectively evaluating and ranking hotspot mutations despite the prevalence of compensatory interactions in allostery systems, our work underscored the value of explicitly computing the functional free energy landscapes to provide quantitative guidance to allosteric modulation for therapeutic and engineering applications.

## Acknowledgement

This work is supported by NIH Grant R35-GM141930. Computational resources from ACCESS (project BIO230101) are greatly appreciated. Part of the computational work was performed on the Shared Computing Cluster which is administered by Boston University’s Research Computing Services (URL: www.bu.edu/tech/support/research/). Anton 2 computer time (award MCB200062P) was provided by the Pittsburgh Supercomputing Center (PSC) through NIH Grant R01GM116961. The Anton 2 machine at PSC was generously made available by D.E. Shaw Research.

## Supporting Information Available

Additional simulation details, results and schematics for a revised thermodynamic model are included.

