## Supplementary Information for "Modulation of Allostery with Multiple Mechanisms by Hotspot Mutations in TetR"

### Additional Computational Methods

#### System Setup and Equilibrium MD Simulations

The initial structure for the ligand-bound simulations is the crystal structure (PDB code: 4AC0) of TetR(B) with bound $Mg^{2+}$ and minocycline. Minocycline is converted to the ligand used in the experimental studies 1, anhydrotetracycline (aTC) treated using the CHARMM General Force Field (CGENFF) model.[2] The protonation state of the ligand is adapted based on the computational analysis of Simonson and co-workers.[3,4] The protein and $Mg^{2+}$ ion are treated using the standard CHARMM force field model.[5] His64, which forms close interaction with the ligand, is protonated at the N$\epsilon$ position.

For the DNA-bound state, the DNA-bound TetR(D) crystal structure (PDB code: 1QPI) is used as the template to first generate a structural model for the DNA-bound TetR(B) using MODELLER[6] (the level of sequence homology between TetR(B) and TetR(D) is 66%). A short targeted MD is then performed to pull the ligand-bound TetR(B) crystal structure (without the ligand-$Mg^{2+}$ complex) towards the modeled DNA-bound TetR(B) conformation. The final structure after 20 ps targeted MD simulation is combined with the DNA 15-mer (sequence CCTATCAATGATAGA) from the DNA-bound TetR(D) crystal structure to establish the initial coordinates of the TetR(B)-DNA complex for subsequent simulations. CHARMM36m[5] is used for both protein and DNA[7] to maintain consistency.

The initial structure of apo TetR(B) is generated in the same way with the apo TetR(D) crystal structures as the template (PDB code: 1BJZ). To facilitate the exploration of the broad conformational space in the apo state, two more initial structures of the apo state are generated by deleting the ligand-$Mg^{2+}$ complex and DNA in the ligand-bound (apo ligand) and DNA-bound (apo dna) TetR(B) structures, respectively.

CHARMM-GUI[8] is used to model the missing protein residues, solvate proteins in TIP3P water[9] cubic boxes with a 10.0 Å edge distance under periodic boundary conditions (PBC), and randomly place 150 mM NaCl ions to neutralize the system and mimic the physiological condition. Particle-mesh Ewald (PME) summation[10] is used to calculate the electrostatic interactions. Van der Waals (vdW) interactions are treated by a cutoff distance of 12 Å and a switch distance of 10 Å. SHAKE algorithm is used in all simulations to constrain bonds involving hydrogen atoms. Systems are energy minimized with steepest descent (SD) and Adopted Basis Newton-Raphson (ABNR) algorithms before equilibration runs with a time step of 1 fs in CHARMM[11] for 500 ps in NVT (constant particle number, volume and temperature) ensemble. Weak harmonic restraints are applied to protein backbone (force constant: 1 kcal $mol^{-1}$ $Å^{-2}$)) and side chain (force constant: 0.1 kcal $mol^{-1}$ $Å^{-2}$)) heavy atoms during equilibration runs. NPT (constant particle number, pressure and temperature) production runs are carried out with a time step of 2 fs in OpenMM 7.3[12] with GPU acceleration. All simulations are maintained at 303.15 K and under 1 bar. Langevin integrator with a friction coefficient of 1 $ps^{-1}$ and Monte Carlo Barostat with the pressure coupling frequency of 100 steps are used in OpenMM production runs.

For the Anton 2 simulations,[13] the equilibration runs are extended to 5 ns in CHARMM. The CHARMM topologies, final coordinates and velocities after equilibration runs are used to convert the system into Anton 2 compatible format in VMD. The Anton 2 production runs are performed in the NPT ensemble. The multigrator scheme[14] is used to allow for the separate updates of the thermostat, barostat, and Newtonian particle dynamics. The simulation temperature and pressure are controlled with a Nosé-Hoover thermostat[15,16] and a Martyna-Tobias-Klein (MTK)[17] barostat with isotropic scaling, respectively. The thermostat and barostat are updated every 24 steps and every 480 steps, respectively. vdW interactions are calculated with a cutoff of 9 Å, and long-rage electrostatic interactions are calculated using the u-series method based on Gaussian split Ewald.[18] The reversible reference system propagator algorithm (RESPA)[19] is used with a time step of 2.4 fs to integrate the long-range non-bonded forces every 5 steps and short-range non-bonded and bonded forces at every step. Production runs are saved every 10 ps and 240 ps in OpenMM and Anton 2 runs respectively.

Overall, independent simulations on local GPUs and on Anton 2 machines are conducted for the ligand-bound, DNA-bound, and apo states of all systems studied in this work. The lengths of trajectories for each system are summarized in Table S1. Both the ligand-$Mg^{2+}$ complex and the DNA are well bound to the protein throughout the corresponding bound state simulations.

#### Critical Ligand Binding Domain (LBD) Order Parameters

LBD order parameters are chosen based on the same Jensen-Shannon (JS) divergence analysis in the previous work.[20] Briefly, protein residues in which at least one heavy atom is within 5 Å of any heavy atoms in the ligand for at least 75% of the simulation time (∼6 $\mu s$) are defined as the ligand-contacting residues. JS divergence of all pair-wise distances among these ligand-contacting residues between the ligand-bound and DNA-bound simulations are calculated. The pairs that exhibit the largest JS divergences are then identified and their distance distributions examined to describe compactness of the ligand-binding pocket. The two pairs that well distinguish the ligand- and DNA-bound ensembles involve Q109 and E147', whose distance characterizes the separation between $\alpha 7$ in one monomer and $\alpha 8$ in the other monomer; and R104 and I134, whose distance characterizes the separation between l6 and $\alpha 8$.

#### Critical DNA Binding Domain (DBD) Order Parameters

For the changes in DBDs, previous structural and MD analyses[21–24] highlighted several possible structural parameters: the center-of-mass separation between the two DBDs (DBD distance), which controls the contacts between the recognition helices ($\alpha 3$) and the DNA; and the angle between $\alpha 4$ during the simulation and the reference orientation in the DNA-bound crystal structure ($\alpha 4$ angle), which describes the pendulum-like motion during the proposed induction mechanism.[22,25] More details for the definitions and computations of these DBD order parameters can be found in our previous study.[20]

#### Kullback-Leibler (KL) Divergence Analysis

Distributions of the DBD order parameters are calculated for the ligand-bound, DNA-bound, and apo states. To compare the similarity of the distributions of the same order parameters or structural properties between different states, or between WT and mutants, Kullback-Leibler (KL) divergence is computed. A larger KL divergence value indicates a larger difference between the two distributions under comparison. KL divergence between two distributions $P$ and $Q$ is defined as follows:

$$D(P||Q) = \int_{-\infty}^{\infty} p(x) \log\left(\frac{p(x)}{q(x)}\right) dx \tag{1}$$

The probability distribution $Q$ can be interpreted as the approximation of the "true" distribution $P$. KL divergence is not symmetric, depending on the choice of $P$ and $Q$. Therefore, the choices of the "true" distribution $P$ and the approximated distribution $Q$ are specified in the corresponding KL divergence tables for clarity. In metadynamics simulations, all mutants sampled a doubly bound basin as discussed in the main text, leading to a bimodal distribution of the DBD distance in the ligand-bound state. To characterize the distribution overlap between the DNA-bound state and the two individual basins of the ligand-bound state in both WT and mutants, each DBD distribution is split into two parts at the distance of 42.5 Å to represent the doubly-bound basin and the ligand only basin, respectively. KL divergence between the DNA-bound distribution and the two individual parts in the ligand-bound state are computed separately.

#### Principal Component Analysis

Principal component analysis (PCA) is a dimensionality reduction technique. Correlated motions of protein residues are characterized by a 3N × 3N covariance matrix of C$\alpha$ coordinates:

$$\mathbf{C} = \left\langle (\mathbf{q} - \langle\mathbf{q}\rangle)(\mathbf{q} - \langle\mathbf{q}\rangle)^T \right\rangle \tag{2}$$

where N is the number of atoms used in PCA and $\mathbf{q} = (x_1, y_1, z_1, x_2, y_2, z_2, \ldots, x_N, y_N, z_N)^T$. After normalization, the covariance $C_{ij}$ ranges from -1 to 1.

3N eigenvalues, which are the variances of the collective motion along the corresponding eigenvectors (i.e. principal components, PCs) sorted in the descending order, are obtained by diagonalization of the 3N × 3N covariance matrix without any filtering. PCs with large variances capture the large magnitude collective motions and are often used to describe the complex motions of the biomolecular system of interest.[26] To illustrate the motions of the DBDs in the apo state simulations, the first principal component (PC1) is chosen as one collective variable of the 2D free energy landscapes for the apo states.

Although only the apo state trajectories are projected onto the PC1, PCs are obtained using the combined trajectories of the ligand-bound, DNA-bound, and apo states of the WT and all mutants with the coordinates of C$\alpha$ atoms for consistency. Inclusion of bound state trajectories also helps define the end states. All frames are aligned against the crystal structure of the WT protein to keep the same direction of principal components among different TetR mutants.

#### Free Energy Landscape Projection

Two-dimensional (2D) free energy landscapes are generated by first constructing a 2D distribution along the selected collective variables and then converting the probability distribution to free energy by

$$\Delta F(V_1, V_2) = -\left[\ln p(V_1, V_2) - \ln p_{max}(V_1, V_2)\right], \quad (3)$$

where $p(V_1, V_2)$ is an estimate of the joint probability density function based on the 2D histogram of the data $(V_1, V_2)$. To ensure that $\Delta F = 0$ at the free energy minimum, $\ln p_{max}(V_1, V_2)$ is subtracted from the free energy.

One-dimensional (1D) free energy landscape or PMF is also constructed by

$$\Delta F(V_1) = -\left[\ln p(V_1) - \ln p_{max}(V_1)\right] \quad (4)$$

Note that the above equations calculate the reduced free energy difference, which has the unit of $k_B T$ where $k_B$ is the Boltzmann constant and $T$ is the temperature in the unit of Kelvin. For each system, the free energy landscape of the apo state is obtained by collecting statistics from independent trajectories with different initial structures.

#### Contact probability calculations

A contact between two protein residues is formed if the distance between the two $\alpha$ carbon (C$\alpha$) atoms is within 5 Å in a frame. A contact matrix for a given frame consists of elements of 1 and 0. 1 indicates a contact and 0 otherwise. Contact probability matrices of WT, G102D, and R49A in the doubly-bound state are computed by reweighting the corresponding contact matrices from metadynamics simulations. The differences in the probability between a mutant and the WT are shown in the 3D protein structure.

#### Covariance and dynamical network analysis

The Pearson correlation coefficients between the Cartesian coordinates of C$\alpha$ atoms are computed in the same way as our previous study.[1] Dynamical network analysis for the apo state is constructed with the Bio3D package[27] based on the Pearson correlation coefficients. Self-correlations and correlations between nearest-neighbors (i.e. $i$+1 and $i$+2 neighbors) are excluded. Nodes in the network represent protein residues and are connected by edges. The weight ("length") of the edge between two nodes $i$ and $j$ is $w_{ij} = -\ln c_{ij}$, where $c_{ij}$ is the Pearson correlation coefficient between $i$ and $j$. The shortest path between two nodes is found using the Floyd-Warshall shortest path algorithm, which minimizes the total length between two nodes. The source residues are the DNA-binding residues (residue ID 2 to 47) and the sink residues are the ligand-contacting residues. 10 suboptimal paths including the shortest path are found for each pair of source and sink residues. In each suboptimal path, residues between the source and sink residues are recorded and the occurrences of these residues are counted. Residues with high occurrences ($> 40\%$) are shown in either the 3D protein structure or a 2D heatmap. Distributions of the path lengths are also plotted.

#### Additional discussion of dynamic network analysis

We conduct network analysis including suboptimal path calculations, which represent a widely used strategy to characterize communication pathways and hub residues that are likely relevant to protein allostery.[28–33] We focus on the apo state for the hotspot mutants. Our previous analysis of the WT protein suggested that motional correlations are stronger and longer-ranged than those in the bound states, which are rigidified by ligand/DNA binding.[20] The distributions of the path-lengths from the sub-optimal path analysis are shown in Fig. S14 and compared using violin plots in Fig. S15. It has been suggested that the path-length distribution reflects the strength of coupling between the allosteric and active sites,[29–32] although the current results do not appear to exhibit any compelling trends in this regard. While some hotspot mutations lead to substantially broader path-length distributions than the WT (e.g., V145A and L127D), several hotspot mutations, especially G102D, lead to notably narrower distributions with lower average values. Since all the mutants studied here exhibit the non-inducible phenotype, our observation suggests that path-length distribution is not well correlated with the induction function.

Another quantity of interest in sub-optimal path calculations is the occurrence, which quantifies the probability that each residue is involved among the shortest paths that connect the allosteric and active sites; residues with high occurrences are often referred to as hub residues and considered to be potentially important to the allosteric coupling.[29–32] We compare the hub residues with high occurrences (>40%) among the mutants in Fig. S16 and map these onto the protein structure in Fig. S17. In the WT, hub residues are observed in the DBDs, loop $l6$, $\alpha7$ and the middle region of $\alpha8$. Some hotspot mutations significantly reduce the number of high-occurrence residues compared to the WT. In G102D, for example, hub residues are no longer observed in $l6$ and $\alpha8$, while in L127D, hub residues are not observed in the DBDs. On the other hand, in R49A, there are many more residues with high occurrences compared to the WT. Therefore, while it is clear that hotspot mutations modify the coupling pathways and therefore the location and density of hub residues, it is difficult to draw conclusion regarding the function of the hotspot mutants based on the hub residue analysis.

Table S1: Summary of Unbiased MD Simulations

| | WT | G102D | R49A | G138H | L127D | G138D | V145A | I193A | G196D |
|---|---|---|---|---|---|---|---|---|---|
| Ligand-bound state | 1 $\mu s$ + 5 $\mu s$ | 1 $\mu s$ + 5 $\mu s$ | 1 $\mu s$ + 5 $\mu s$ | 1 $\mu s$ + 5 $\mu s$ | 1 $\mu s$ + 5 $\mu s$ | 1 $\mu s$ + 5 $\mu s$ | 1 $\mu s$ + - $\mu s$ | 1 $\mu s$ + 5 $\mu s$ | 1 $\mu s$ + 5 $\mu s$ |
| DNA-bound state | 1 $\mu s$ + 5 $\mu s$ | 1 $\mu s$ + 5 $\mu s$ | 1 $\mu s$ + 5 $\mu s$ | 1 $\mu s$ + 5 $\mu s$ | 1 $\mu s$ + 5 $\mu s$ | 1 $\mu s$ + 5 $\mu s$ | 1 $\mu s$ + - $\mu s$ | 1 $\mu s$ + 5 $\mu s$ | 1 $\mu s$ + 5 $\mu s$ |
| Apo state | 3 $\mu s$ + 50 $\mu s$ | 3 $\mu s$ + 30 $\mu s$ | - $\mu s$ + 30 $\mu s$ | - $\mu s$ + 30 $\mu s$ | 9 $\mu s$ + 30 $\mu s$ | 9 $\mu s$ + 30 $\mu s$ | 9 $\mu s$ + - $\mu s$ | 9 $\mu s$ + 30 $\mu s$ | - $\mu s$ + 30 $\mu s$ |

a. The lengths of trajectories are listed in the format of "local GPUs + Anton 2". "-" indicates that simulation is not carried out on local GPUs or Anton 2.

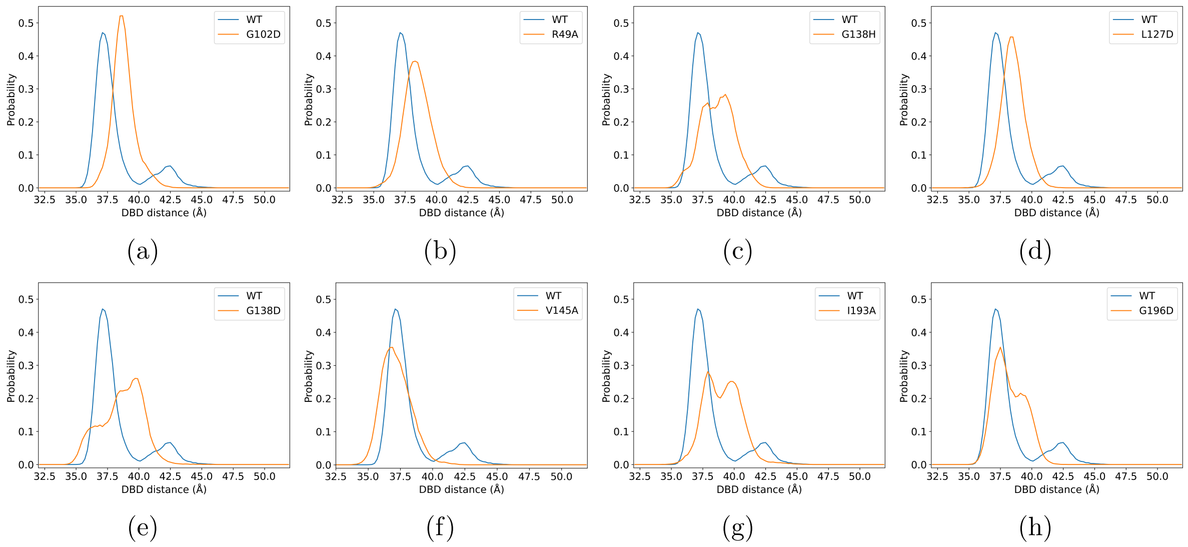


Figure S1: The comparison of the DBD distance distributions for the DNA-bound states between WT and hotspot mutants based on unbiased MD simulations. (a) G102D, (b) R49A, (c) G138H, (d) L127D, (e) G138D, (f) V145A, (g) I193A, and (h) G196D. See Table S2 for the KL divergence values.

Table S2: The Kullback-Leibler divergence analysis for the DBD distance distributions: the comparison between WT and mutant DNA-bound states based on unbiased MD simulations.

| G102D | R49A | G138H | L127D | G138D | V145A | I193A | G196D |
|---|---|---|---|---|---|---|---|
| 1.711 | 1.121 | 1.243 | 1.297 | 1.804 | 1.239 | 1.404 | 0.752 |

a. The reference is the WT distribution. See Fig. S1 for the distributions.

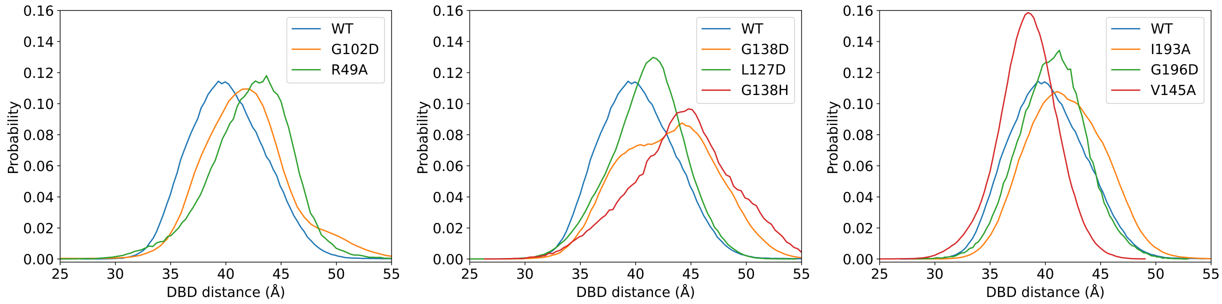


(a) (b) (c)

Figure S2: The comparison of the DBD distance distributions for the apo states between WT and mutants based on unbiased MD simulations.

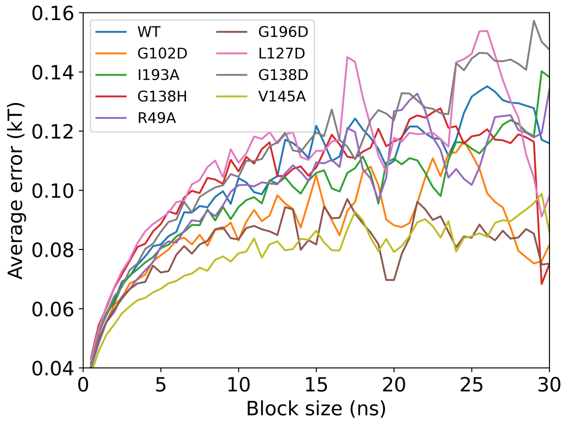


Figure S3: Statistical error analysis for the convergence of PMF calculations based on metadynamics simulations.

Table S3: The Kullback-Leibler divergence analysis for the $\alpha 4$ angle distributions: the comparison between ligand and DNA-bound states based on unbiased MD simulations.

| | WT | G102D | R49A | G138H | L127D | G138D | V145A | I193A | G196D |
|---|---|---|---|---|---|---|---|---|---|
| $\alpha 4$ angle | 46.803 | 22.756 | 25.846 | 6.207 | 27.578 | 13.196 | 52.554 | 4.172 | 33.614 |

a. The reference is the ligand-bound state distribution. See Fig. S6 for the distributions.

Table S4: The Kullback-Leibler divergence analysis for the distributions of l6-$\alpha 8$ distance in the ligand binding pocket (C$\alpha$ distance between R104 and L134): comparison of ligand only basin and doubly bound basin based on metadynamics simulations.

| WT | G102D | R49A | G138H | L127D | G138D | V145A | I193A | G196D |
|---|---|---|---|---|---|---|---|---|
| 0.038 | 0.260 | 0.019 | 0.533 | 0.053 | 0.186 | 0.200 | 0.162 | 0.125 |

a. The reference is the distribution in the ligand-only basin of each system. See Fig. S10 for the distributions.

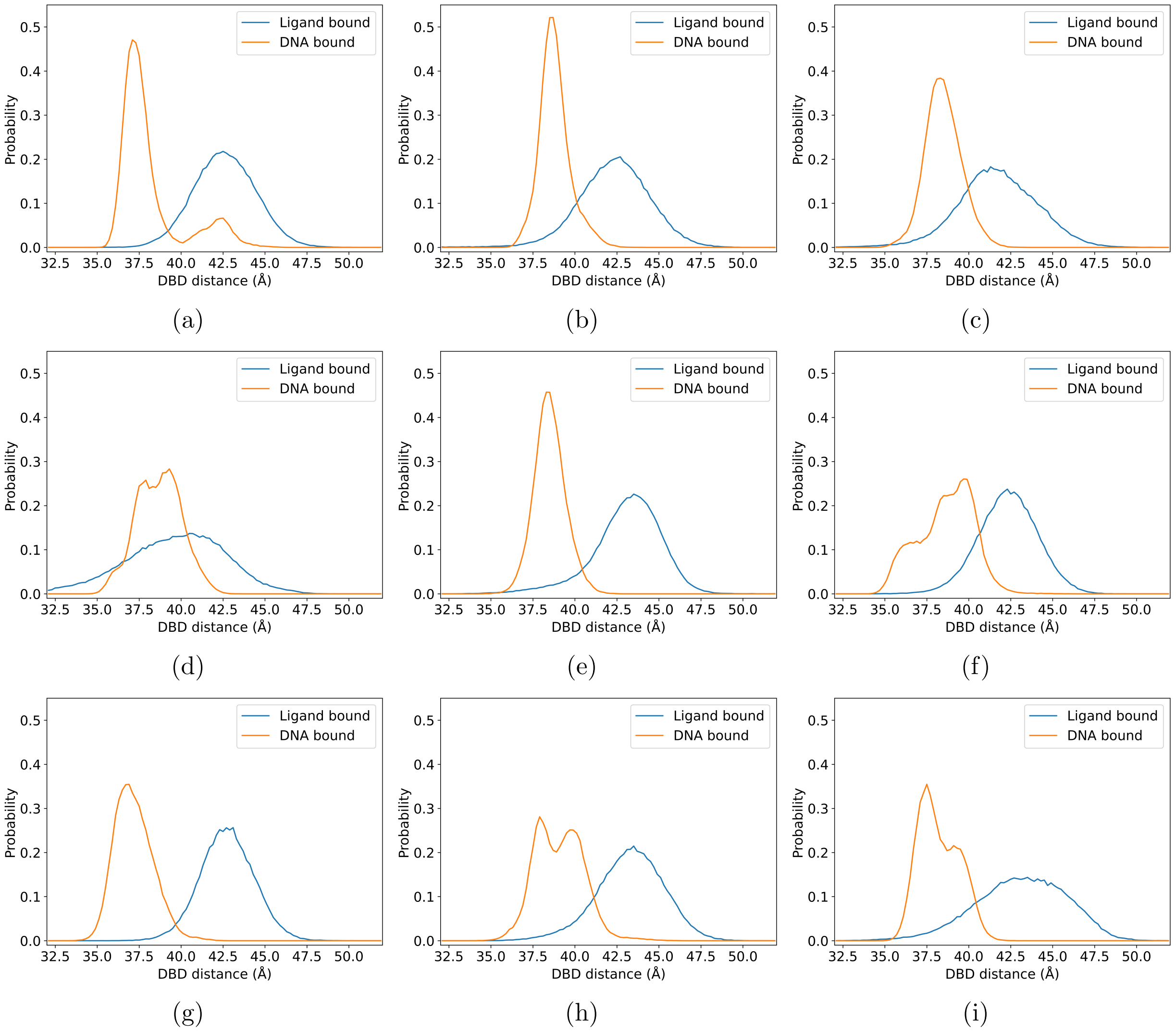


Figure S4: The comparison of the DBD distance distributions between ligand and DNA-bound states based on unbiased MD simulations. (a) WT, (b) G102D, (c) R49A, (d) G138H, (e) L127D, (f) G138D, (g) V145A, (h) I193A, and (i) G196D.

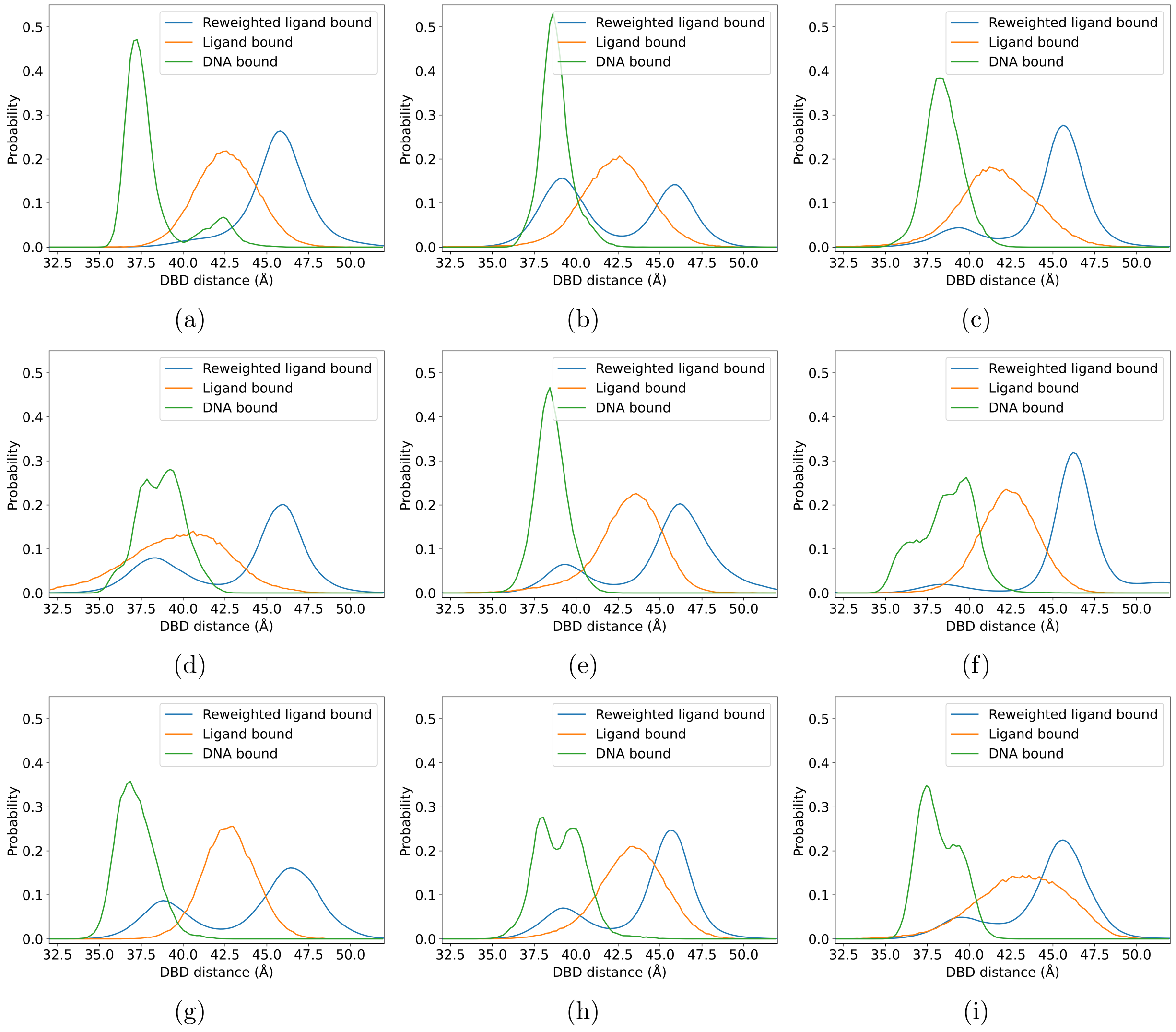


Figure S5: The same as Fig. S4, with the additional reweighted distribution from metadynamics simulations for the ligand-bound states.

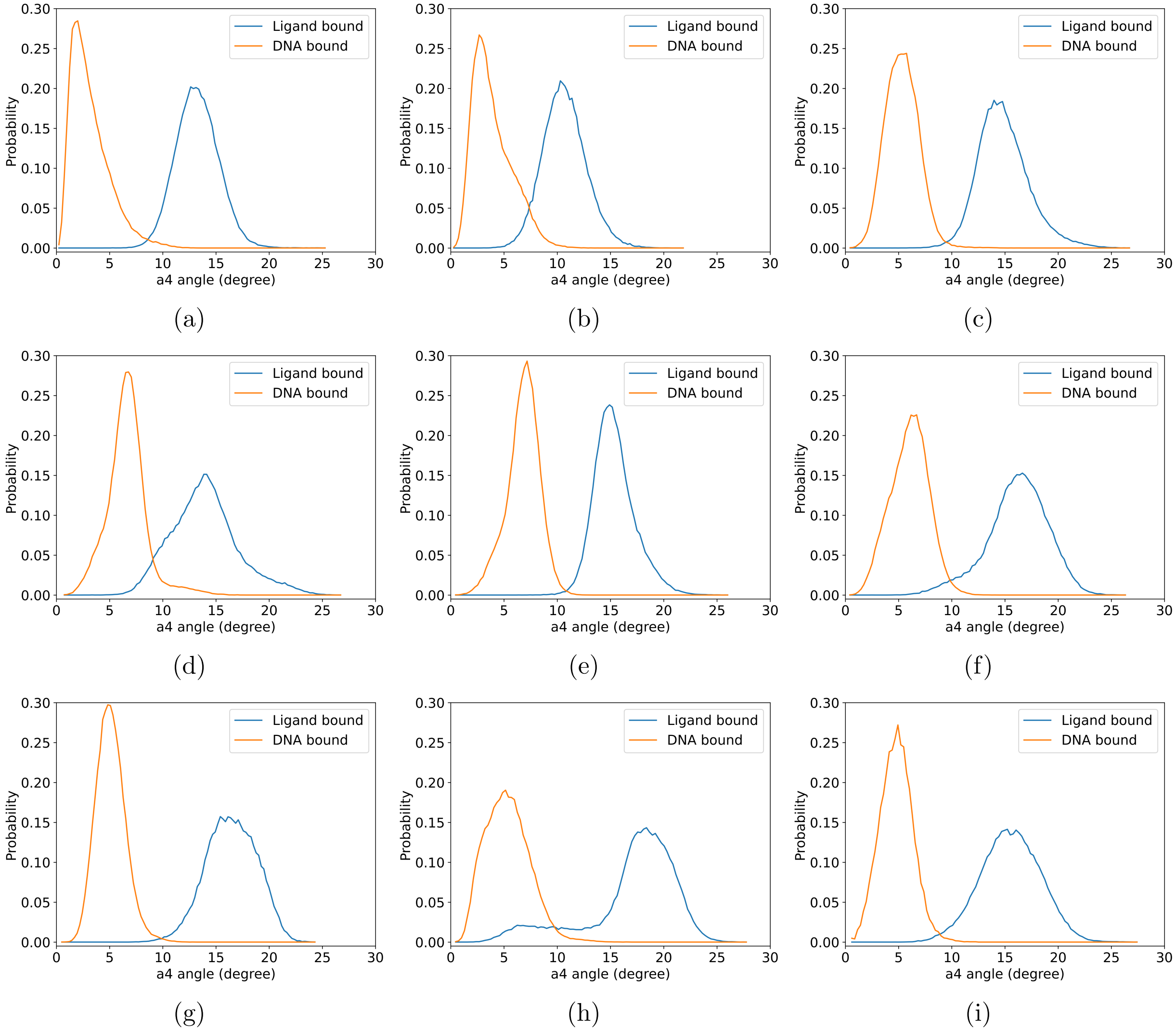


Figure S6: The comparison of the $\alpha 4$ angle distributions between ligand and DNA-bound states based on unbiased MD simulations. (a) WT, (b) G102D, (c) R49A, (d) G138H, (e) L127D, (f) G138D, (g) V145A, (h) I193A, and (i) G196D. See Table S3 for the KL divergence values.

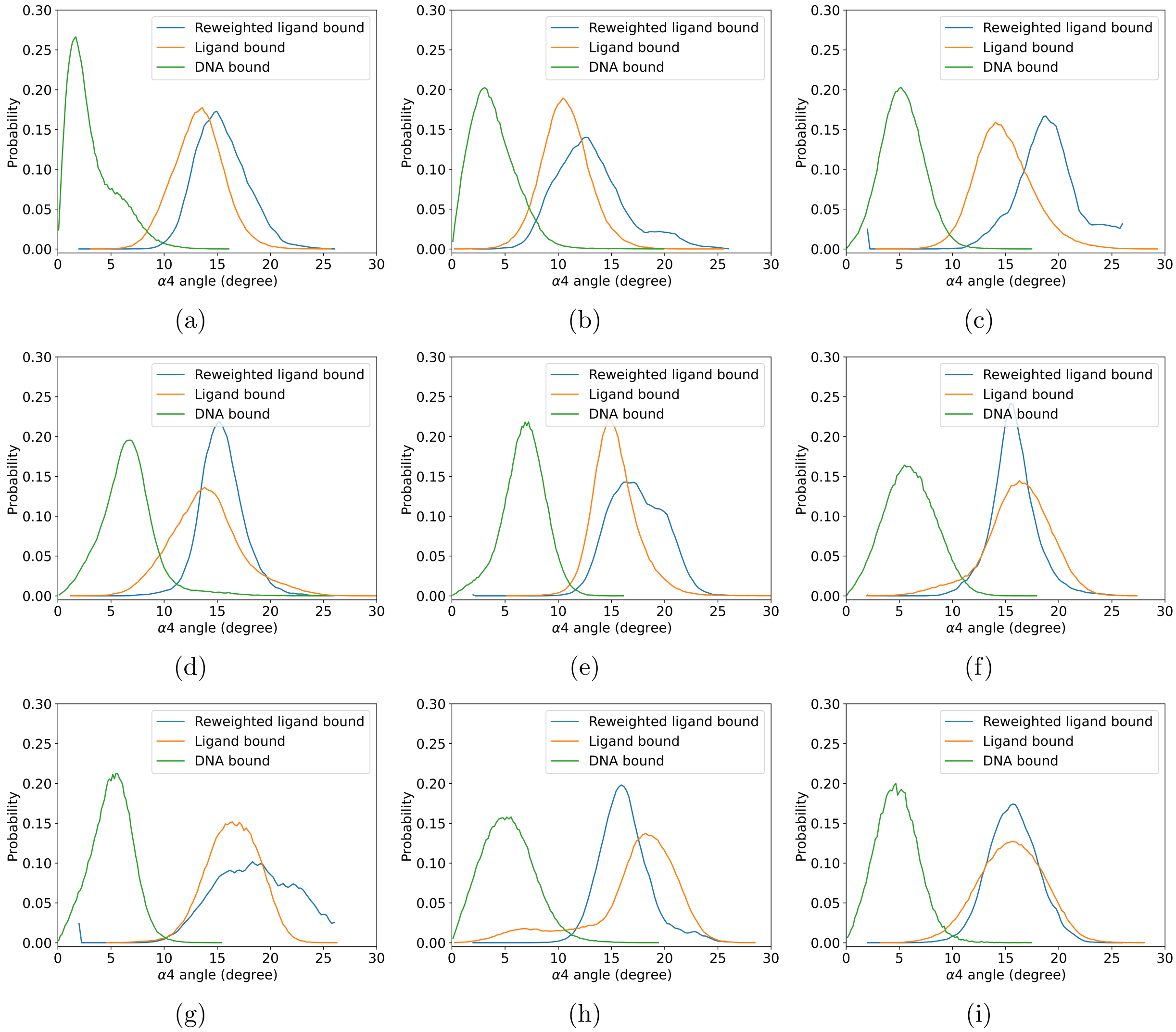


Figure S7: The same as Figure S6, with the additional reweighted distributions from metadynamics simulations for the ligand-bound states.

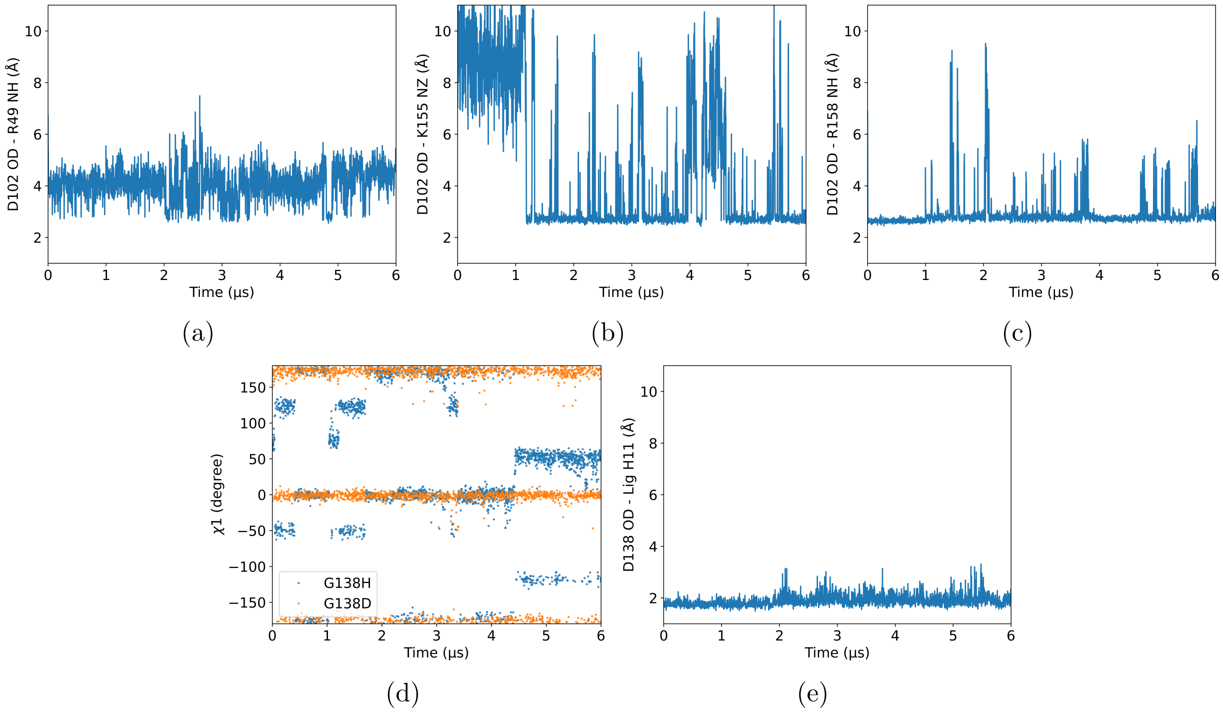


Figure S8: Key structural properties that characterize the local environment of several mutated residues during the unbiased MD simulations of the ligand-bound states. (a-c) D102 in G102D engages in interactions with the cationic residues in $\alpha4$ and $\alpha8$': R49, K155' and R158', respectively. Distances are the minimal distances between OD 1/2 of D102 and NH 1/2 of R49, NH 1/2 of R158', or NZ of K155'. (d) Evolution of the $\chi_1$ angle of residue 138 in G138H and G138D. (e) Evolution of hydrogen bonding distances between OD 1/2 of D138 and H11 of the ligand in the G138D simulation.

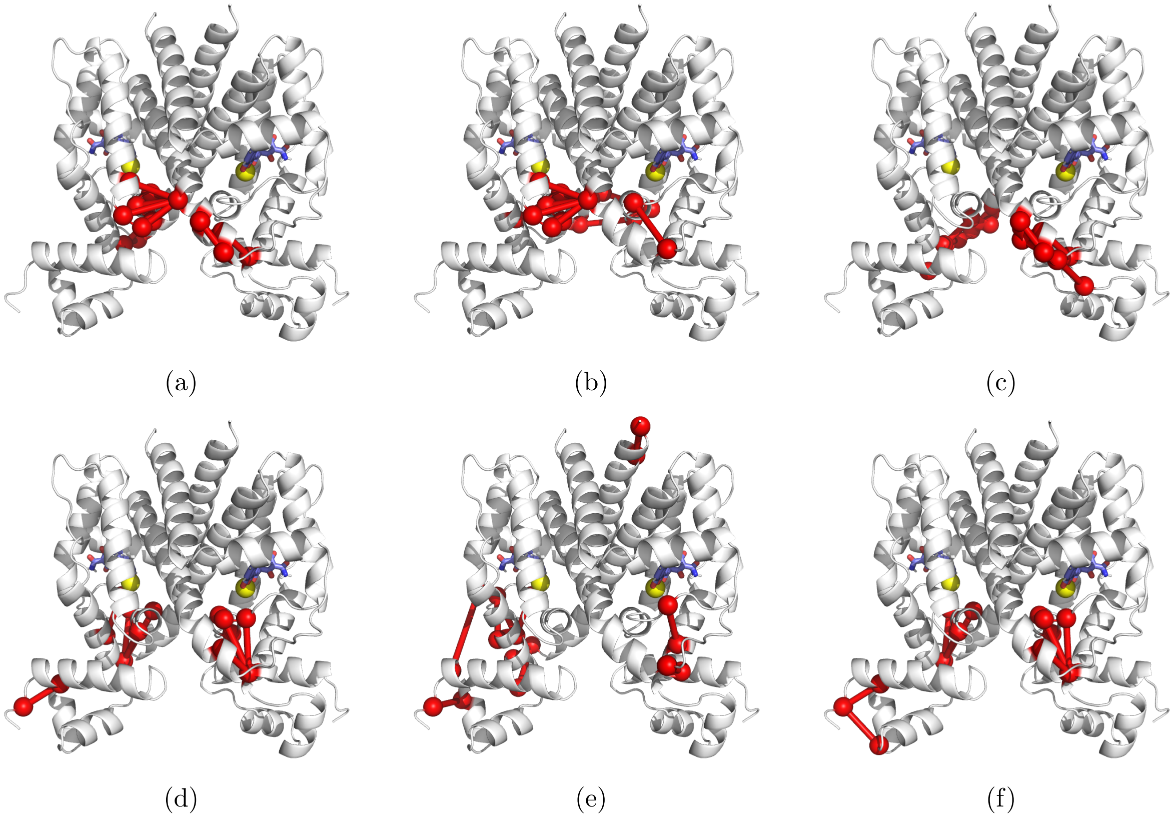

Figure S9: Residues that undergo large changes in contact probability relative to the WT in the doubly-bound basin. The residues are mapped onto the protein structure with the contacts indicated as bonds. The first row is for G102D, and the second row for R49A: (a and d) top ten largest absolute change, (b and e) top ten increase and (c and f) top ten decrease in contact probabilities relative to the WT, respectively. The ligands are shown in licorice, and the $Mg^{2+}$ ions are shown as yellow spheres.

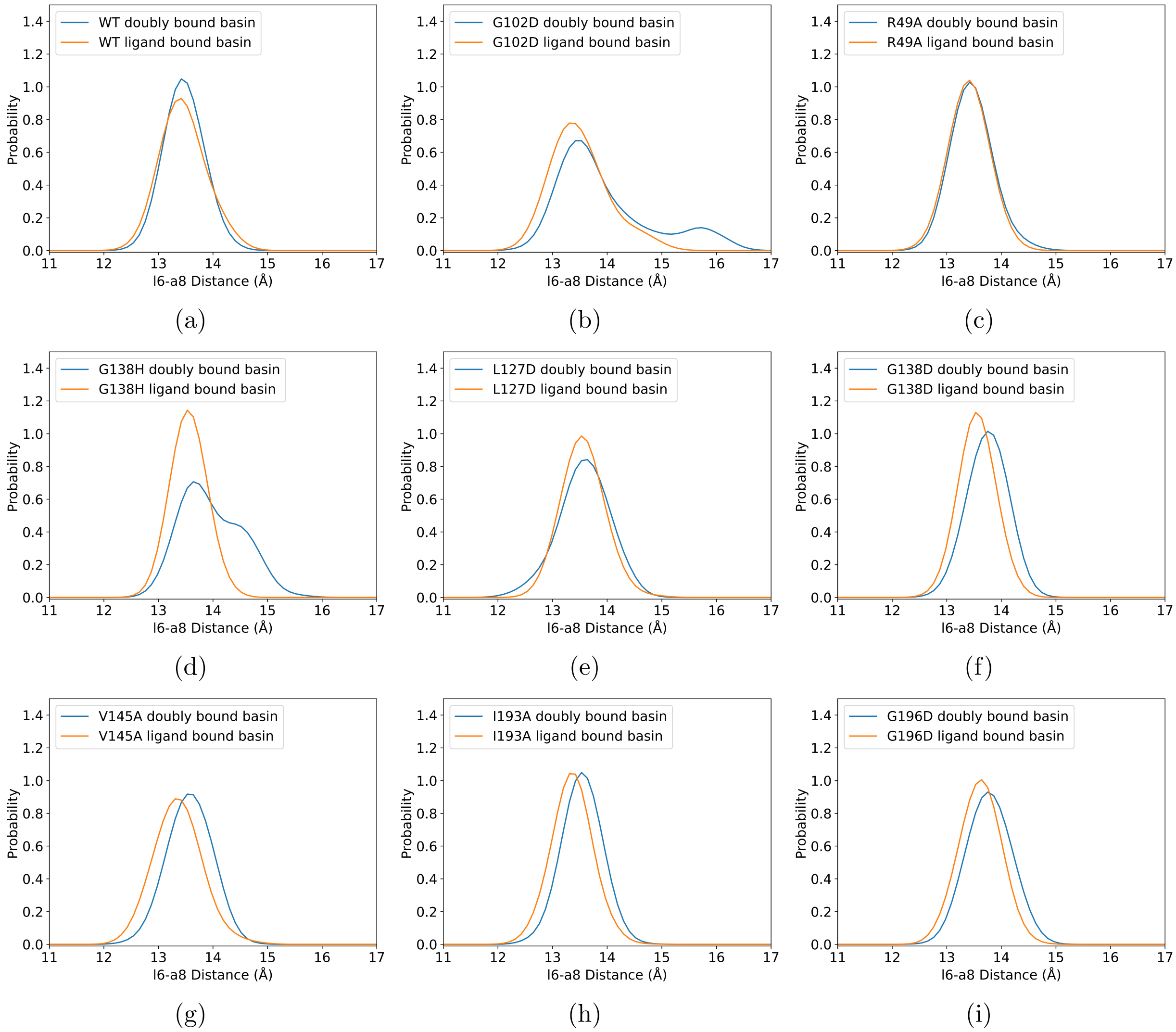


Figure S10: The reweighted distributions of the distance between R104 and L134 (l6-$\alpha$8) in the "doubly-bound basin" (DBD distance range is between 37.5 Å and 38.0 Å) and the "ligand-only basin" (DBD distance range is between 46.5 Å and 47.0 Å) based on metadynamics simulations in the ligand-bound states. See Table S4 for the KL divergence values.

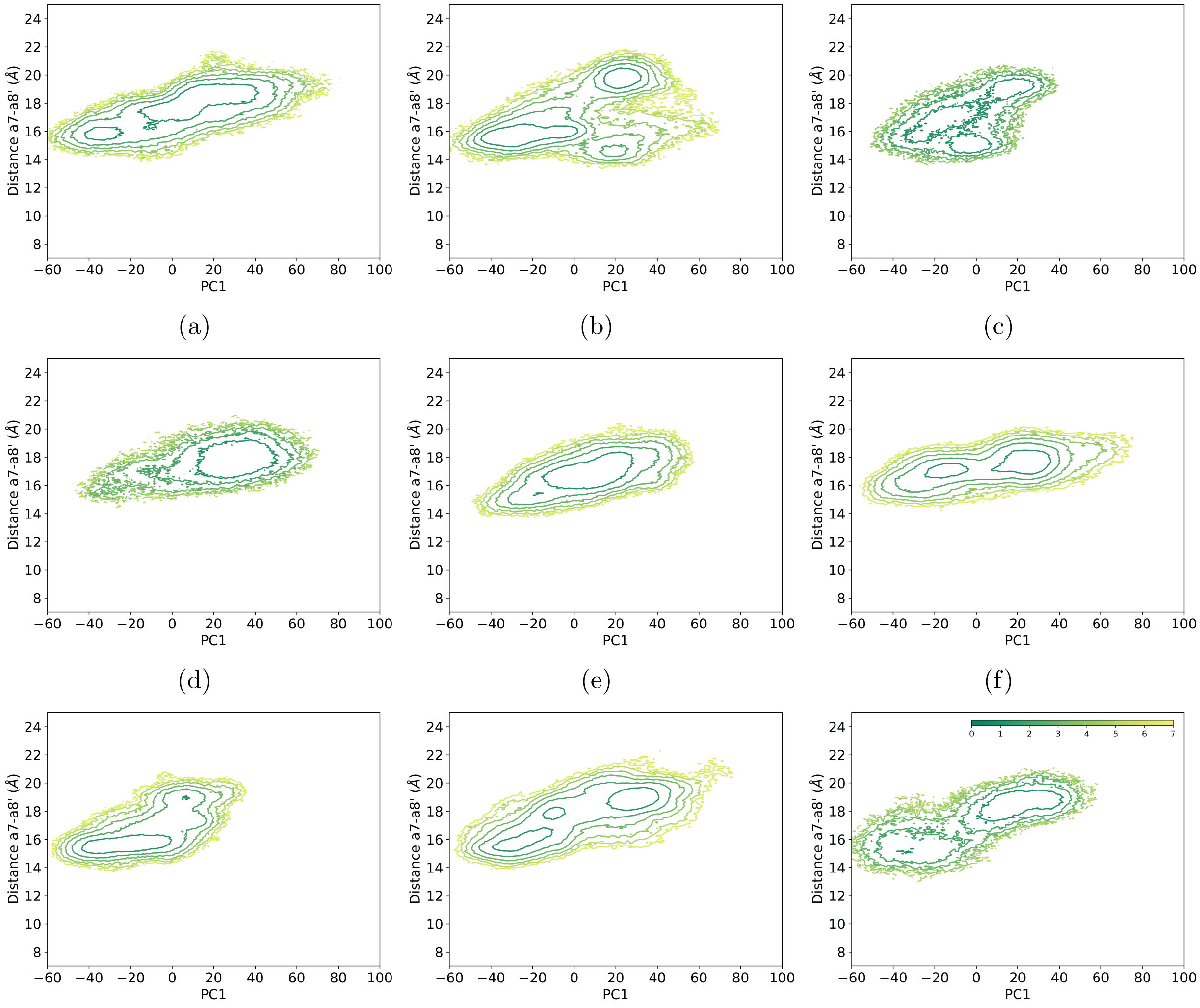


(g) (h) (i)

Figure S11: Two-dimensional free energy landscapes of the apo state spanned by the first principal component (PC1) and $\alpha7-\alpha8'$ distance (C$\alpha$ distance between Q109 and E147'). (a) WT, (b) G102D, (c) R49A, (d) G138H, (e) L127D, (f) G138D, (g) V145A, (h) I193A, and (i) G196D. Color bar is in the unit of $k_B$T.

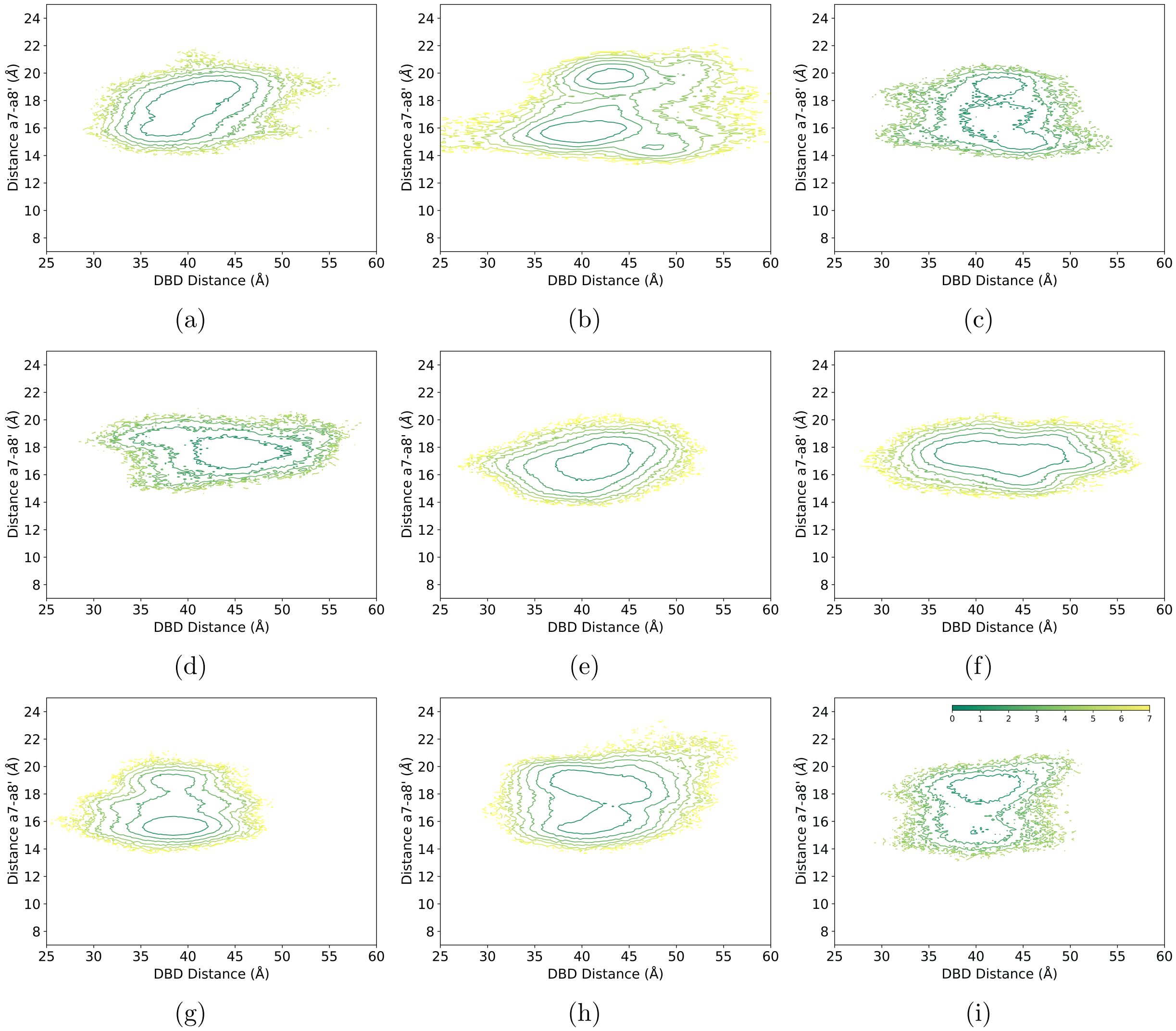


Figure S12: Two-dimensional free energy landscapes of the apo state spanned by the DBD distance and $\alpha7-\alpha8'$ distance (C$\alpha$ distance between Q109 and E147'). (a) WT, (b) G102D, (c) R49A, (d) G138H, (e) L127D, (f) G138D, (g) V145A, (h) I193A, and (i) G196D. Color bar is in the unit of $k_B$T.

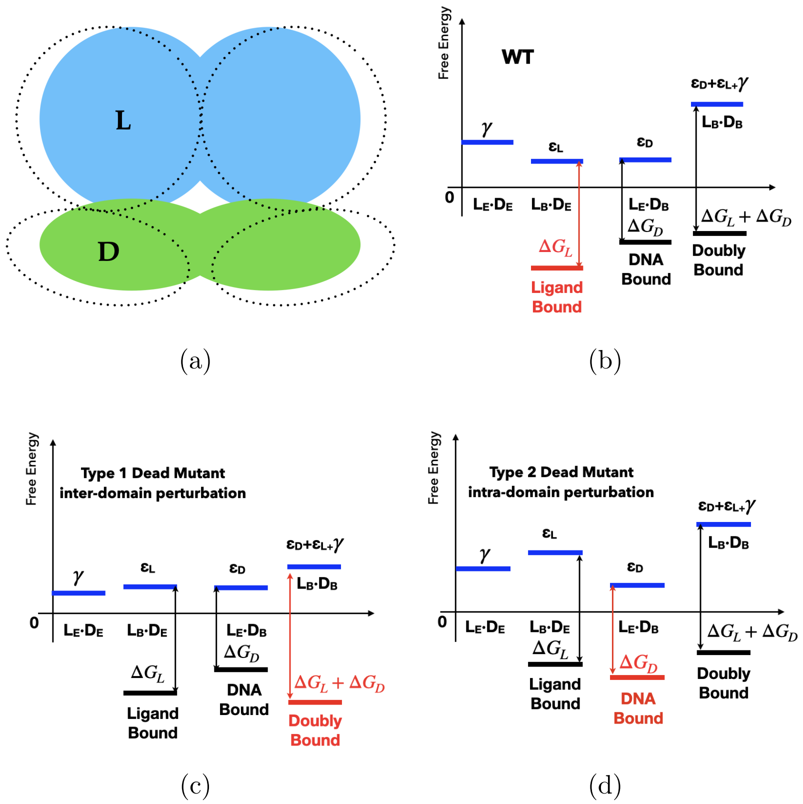


Figure S13: A revised MWC-like model that qualitatively distinguishes two types of allostery hotspots in TetR. (a) Same as our previous work,[20] TetR is divided into LBD (L) and DBD (D), and each can adopt two conformations: the relaxed/inactive conformation is incapable of binding, while the active conformation is binding competent (with intrinsic binding free energies of $\Delta G_L$ and $\Delta G_D$, respectively) but lies at a higher free energy ($\epsilon_L$ for L and $\epsilon_D$ for D). While the original model assumed that an unfavorable coupling free energy $\gamma$ applies only when both domains adopt the active conformation, the revised model posits that $\gamma$ contributes as far as both domains adopt the same (active or inactive) conformation. Panels (b)-(d) follow the same format as Fig. 9 in Ref. 20 to illustrate the qualitative free energy diagrams for the WT protein and two limiting types of dead mutants, respectively; the blue bars indicate the free energy levels for the various conformational states of the apo protein. According to the revised model, the apo landscape in the WT is expected to feature mainly two basins that correspond to $L_B \cdot D_E$ and $L_E \cdot D_B$; with type 1 mutation, $L_E \cdot D_E$ is also lower in free energy, thus the apo landscape is expected to feature three basins, as observed in the computed apo landscape for G102D and R49A (Fig. S11 b,c).

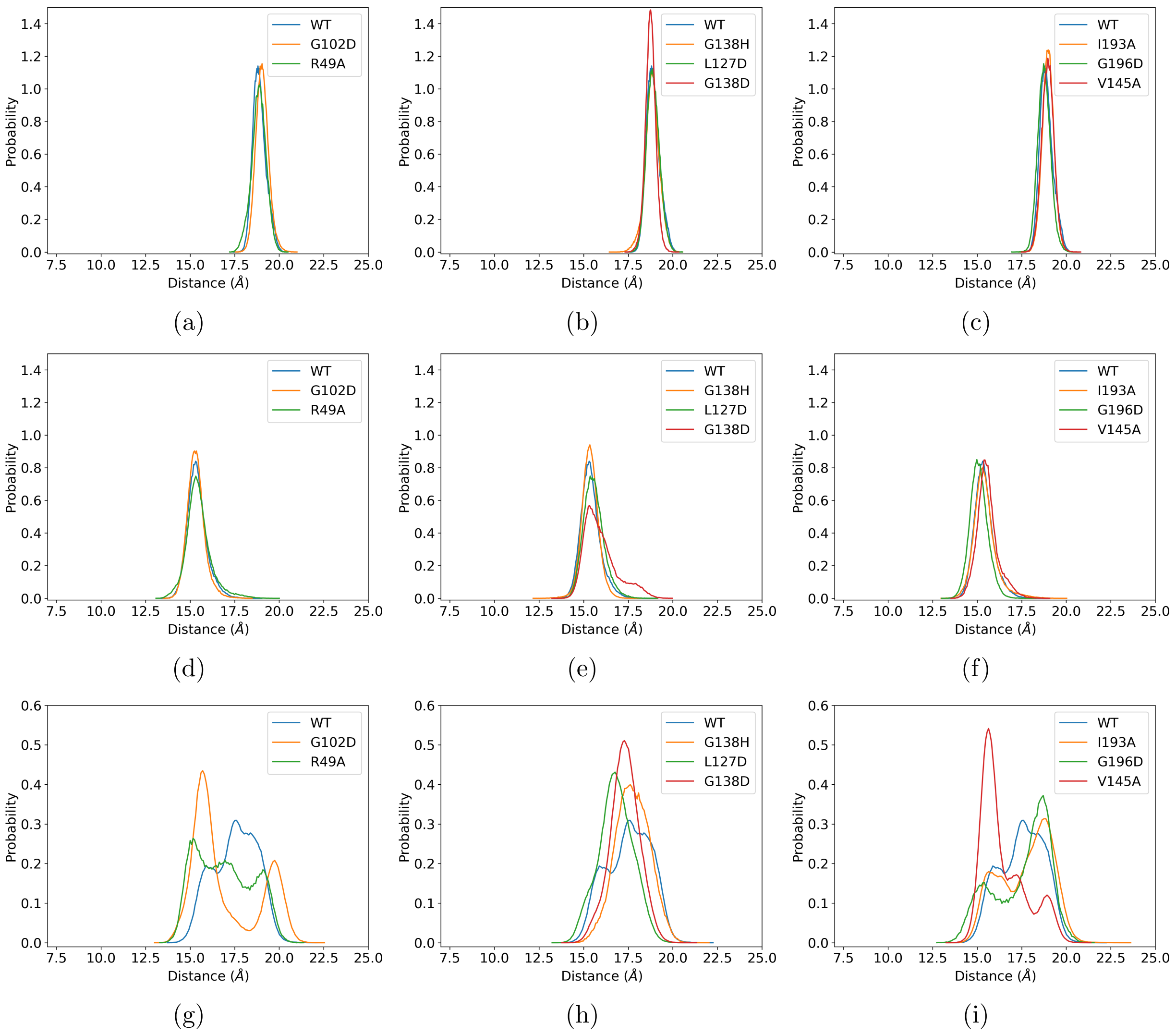


Figure S14: Distributions of $\alpha 7 - \alpha 8'$ distance (C$\alpha$ distance between Q109E and E147') based on unbiased MD simulations. (a-c) Ligand-bound states; (d-f) DNA-bound states; (g-i) apo states.

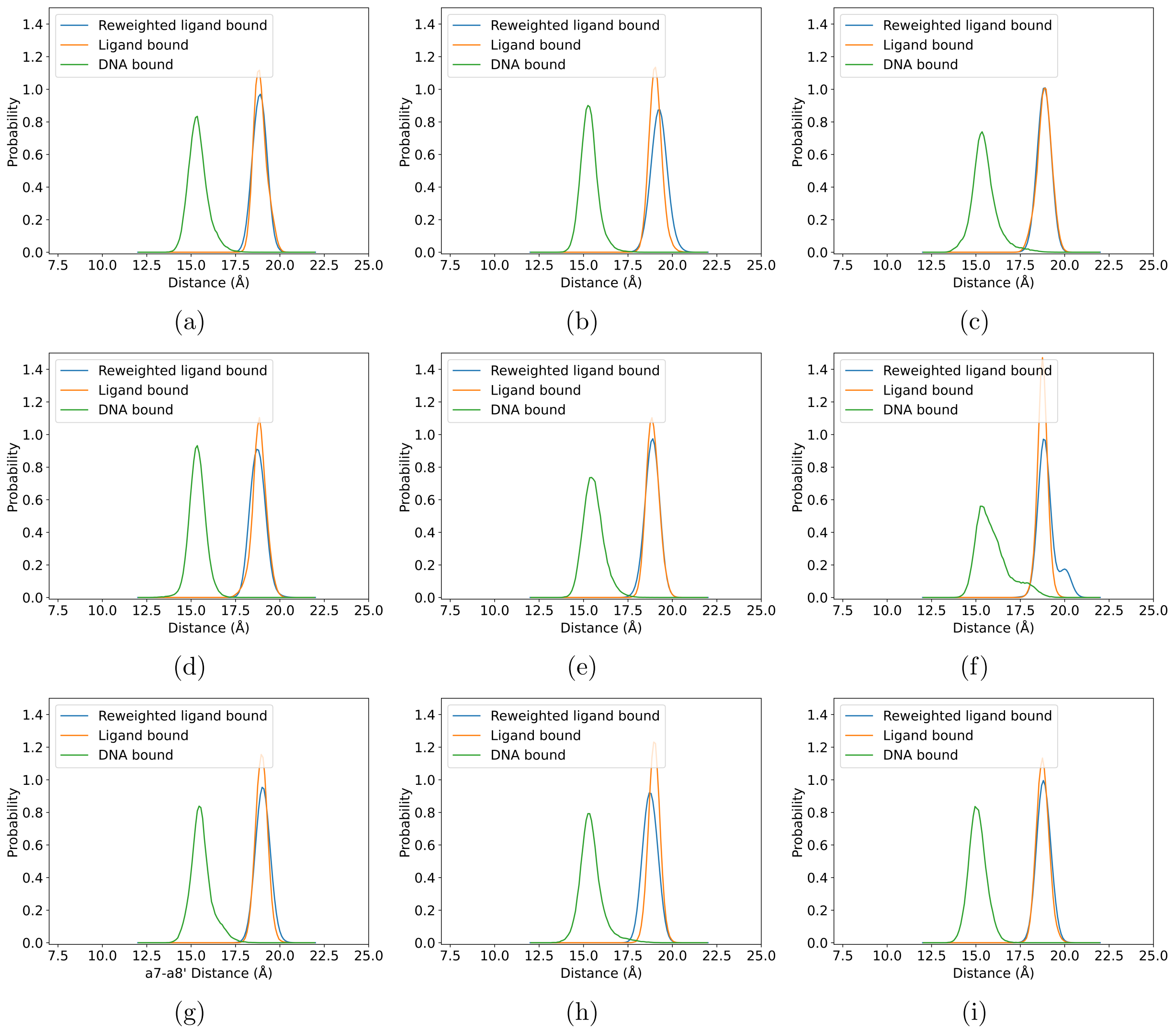


Figure S15: The comparison of $\alpha7 - \alpha8'$ distance distributions (C$\alpha$ distance between Q109E and E147') in the ligand-bound and DNA-bound states based on unbiased MD and metadynamics simulations. (a) WT, (b) G102D, (c) R49A, (d) G138H, (e) L127D, (f) G138D, (g) V145A, (h) I193A, and (i) G196D.

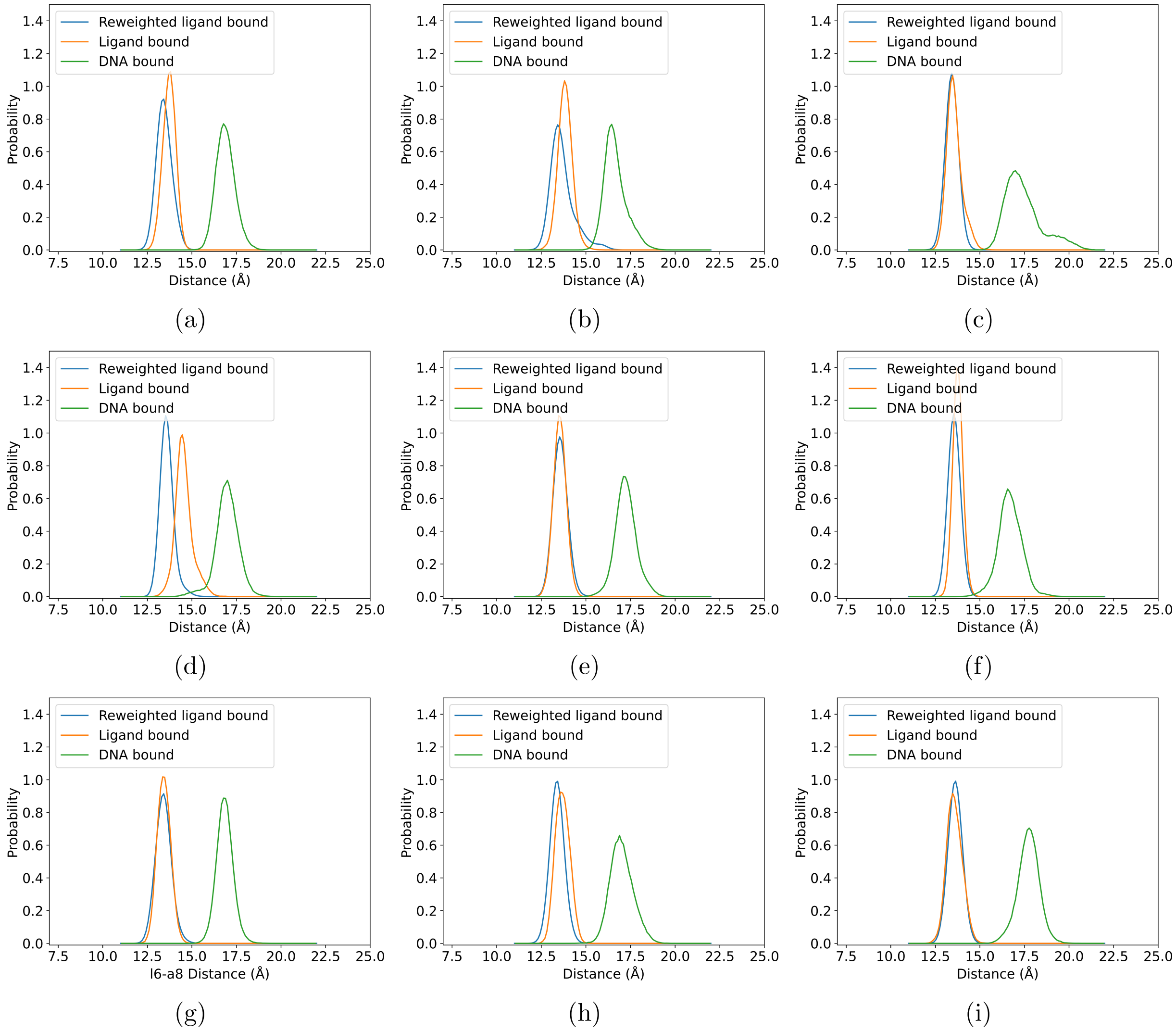


Figure S16: The comparison of l6-$\alpha$8 distance distributions (C$\alpha$ distance between R104 and L134) in the ligand-bound and DNA-bound states based on unbiased MD and metadynamics simulations. (a) WT, (b) G102D, (c) R49A, (d) G138H, (e) L127D, (f) G138D, (g) V145A, (h) I193A, and (i) G196D.

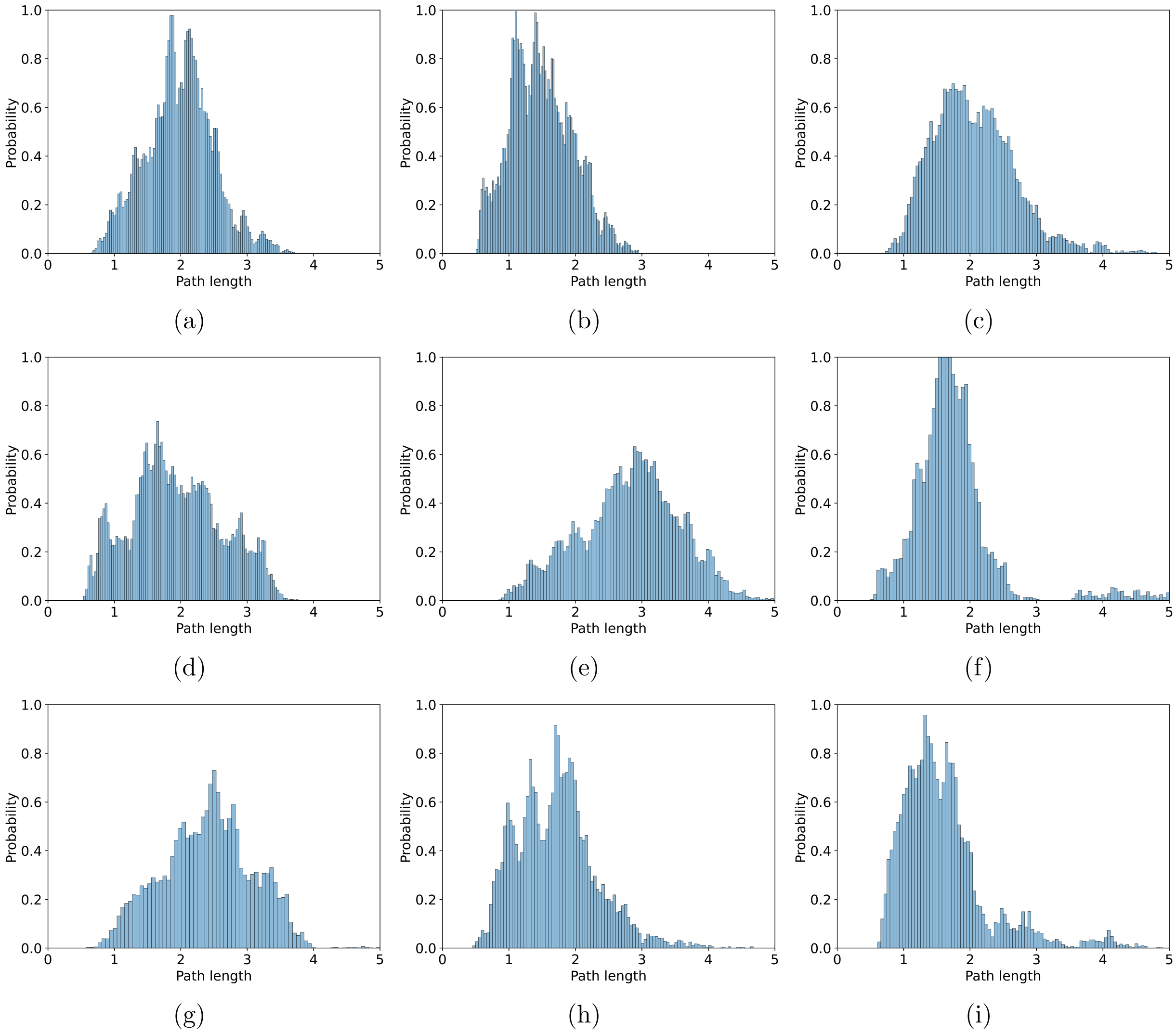


Figure S17: Path-lengths distributions from suboptimal path calculations for the apo state. (a) WT, (b) G102D, (c) R49A, (d) G138H, (e) L127D, (f) G138D, (g) V145A, (h) I193A, and (i) G196D.

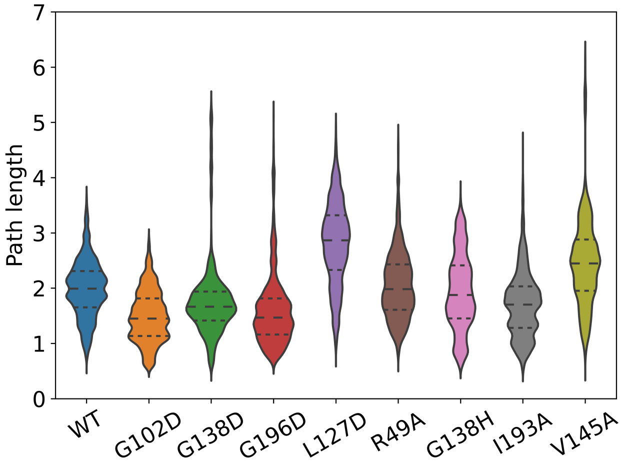


Figure S18: Violin plots for the comparison of path-length distributions (see Fig. S17) from suboptimal path calculations for the apo state of TetR mutants.

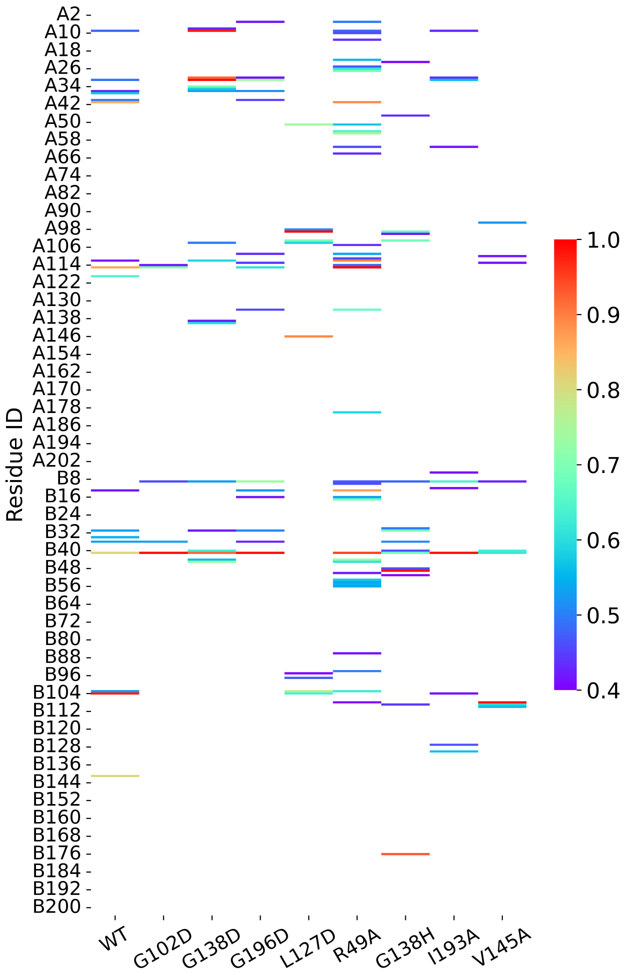


Figure S19: Residues with high occurrences (>40%) from suboptimal path calculations for the apo state of TetR mutants.

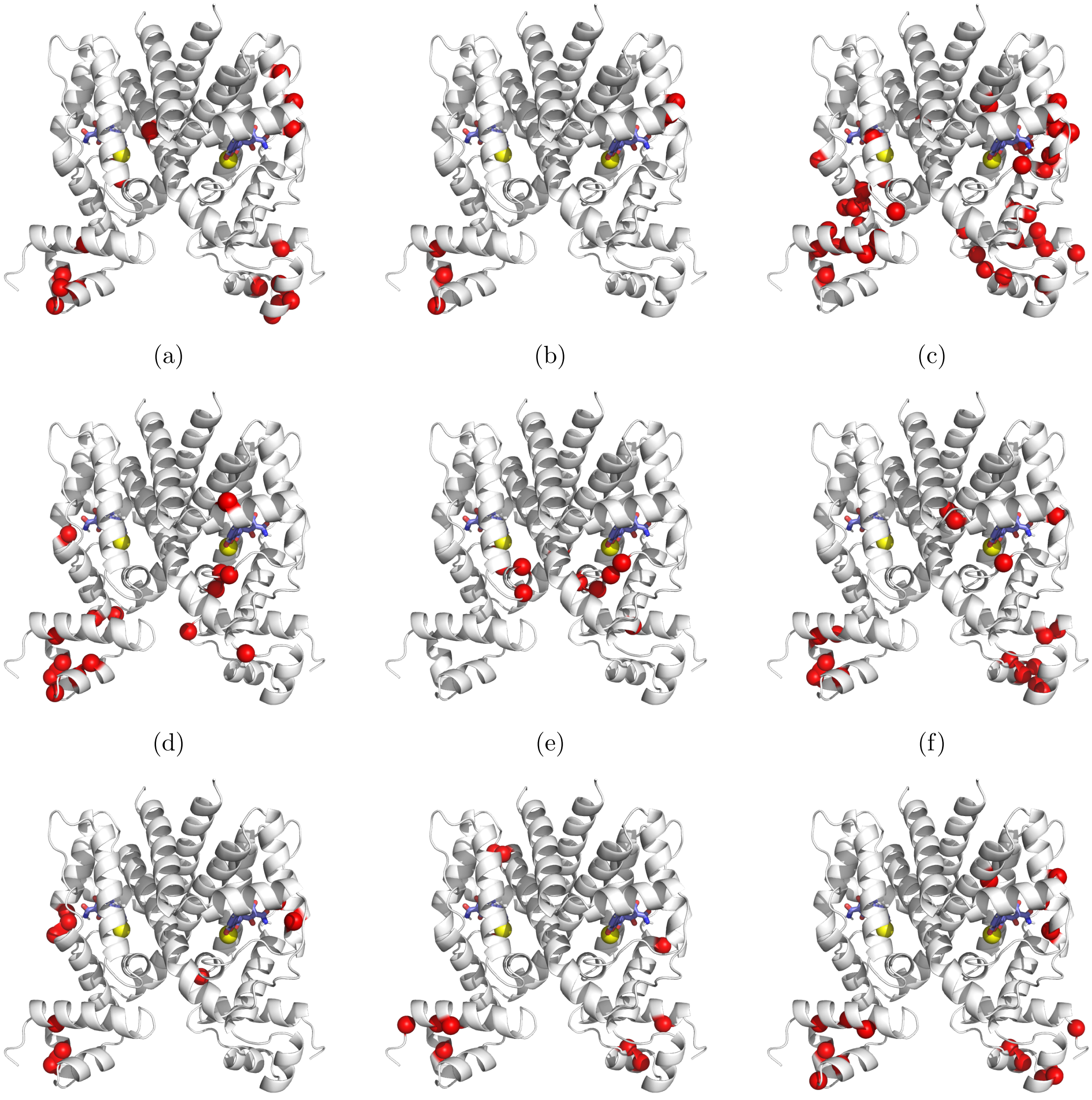

(a) (b) (c)

(d) (e) (f)

(g) (h) (i)

Figure S20: Hub residues (occurrences >40%) from suboptimal path calculations for the apo state of TetR mutants mapped onto the protein structure. (a) WT, (b) G102D, (c) R49A, (d) G138H, (e) L127D, (f) G138D, (g) V145A, (h) I193A, and (i) G196D. The ligands are shown in licorice, and the $Mg^{2+}$ ions are shown as yellow spheres.

(18) Predescu, C.; Lerer, A. K.; Lippert, R. A.; Towles, B.; Grossman, J. P.; Shaw, D. E.

The u-series: A separable decomposition for electrostatics computation with improved accuracy. *J. Chem. Phys.* **2020**, *152*, 084113.
